# Deciphering the contribution of oriens-lacunosum/moleculare (OLM) cells to intrinsic theta rhythms using biophysical local field potential (LFP) models

**DOI:** 10.1101/246561

**Authors:** Alexandra P. Chatzikalymniou, Frances K. Skinner

## Abstract

Oscillations in local field potentials (LFPs) commonly occur and analyses of them fuel brain function hypotheses. An understanding of the cellular correlates and pathways affecting LFPs is needed but many overlapping pathways *in vivo* makes this difficult to achieve. A prevalent LFP rhythm in the hippocampus is ‘theta’ (3-12 Hz). Theta rhythms emerge intrinsically in an *in vitro* whole hippocampus preparation and thus can be produced by local interactions between interneurons and pyramidal (PYR) cells. Overlapping pathways are much reduced in this preparation making it possible to decipher the contribution of different cell types to LFP generation. We focus on oriens-lacunosum/moleculare (OLM) cells as a major class of interneurons in the hippocampus. They can influence PYR cells through two distinct pathways, (i) by direct inhibition of PYR cell distal dendrites, and (ii) by indirect disinhibition of PYR cell proximal dendrites by inhibiting bistratified cells (BiCs) that target them. We use previous inhibitory network models and build biophysical LFP models using volume conductor theory. We assess the effect of OLM cells to ongoing intrinsic LFP theta rhythms by directly comparing our model LFP features with experiment. We find that robust LFP theta responses adhering to reproducible experimental criteria occur only for particular connectivities between OLM cells and BiCs. Decomposition of the LFP reveals that OLM cell inputs onto the PYR cell regulate robustness of LFP responses without affecting average power and that the robust response depends on co-activation of distal inhibition and basal excitation. We use our models to estimate the spatial extent of the region generating LFP theta rhythms, leading us to predict that about 22,000 PYR cells participate in generating the LFP theta rhythm. Besides allowing us to understand OLM cells’ contributions to intrinsic theta rhythms, our work can drive hypothesis developments of cellular contributions *in vivo*.

**Author Summary:** Oscillatory local field potentials (LFPs) are extracellularly recorded potentials that are widely used to interpret information processing in the brain. For example, theta LFP rhythms (3-12 Hz) are correlated with memory processing and it is known that particular inhibitory cell types control their existence. As such, it is critical for us to appreciate how various cell types contribute to the characteristics of LFP rhythms. A precise biophysical modeling scheme linking activity at the cellular level and the recorded signal has been established. However, it is difficult to assess cellular contributions *in vivo* because of many spatiotemporally overlapping pathways that prevent the unambiguous separation of signals. Using an *in vitro* preparation that exhibits intrinsic theta (3-12 Hz) rhythms and where there is much less overlap, we build biophysical LFP models to explore cell contributions to ongoing intrinsic theta rhythms. We uncover distinct contributions from different cell types and show that robust theta rhythms depend specifically on one of the cell types. We are able to determine this because our LFP models have direct links with experiment and we are able to perform thousands of simulations.

## Introduction

Oscillatory brain activities, as can be observed in EEGs and local field potentials (LFPs), are a ubiquitous feature of brain recordings [1]. These activities are brought about by interacting excitatory and inhibitory networks, and accumulating evidence indicates that rhythms can form part of the neural code by phasically organizing information in brain circuits [2]. The local field potential (LFP) is the low-frequency part (<500 Hz) of the extracellularly recorded potential and is a widely recorded signal in experimental configurations. The LFP originates from transmembrane currents passing through cellular membranes in the vicinity of a recording electrode tip [3] and its biophysical origin is understood in the framework of volume conductor theory [4]. Many sources contribute to the LFP [5] and they depend on the frequency range of the extracellular signal [6]. Slower oscillations (< 50 Hz) are generated by synaptic currents as opposed to higher frequency oscillations (> 90 Hz) which are influenced by phase-modulated spiking activity [6]. The ease of LFP recordings explains why they are a common measure of neural activity. However, determining the sources of LFP output is highly challenging in general, and the contributions of remote and local activities can be non-intuitive [7]. In essence, it is far from clear how to interpret LFP recordings in light of contributions from different cell types and pathways.

The hippocampus exhibits many LFP population activities including theta and gamma rhythms [8, 9]. In particular, the theta rhythm (3-12 Hz) is a prominent rhythm that is correlated with spatial navigation and episodic memory, rapid eye movement sleep and voluntary behaviors [10]. Recently, direct behavioural relevance of theta LFP rhythm phase-coding was demonstrated by delivering perturbations during specific phases of the theta rhythm to preferentially affect encoding or retrieval behaviours [11]. This was done by optogenetically stimulating particular inhibitory cell types in the dorsal CA1 region of the hippocampus. Such exciting studies and several reviews [12–14] make it clear that the specifics of inhibitory cell types are fundamental to neural coding and brain function. In essence, if we are to understand the brain’s code, i.e., behaviour-related changes in oscillatory activity, we need to determine and understand how various cell type populations contribute to LFP recordings.

It has long been known that input from the medial septum is an important contributor to *in vivo* LFP theta rhythms [10]. However, recent work by Goutagny and colleagues showed that theta rhythms can emerge in the CA1 region of an intact *in vitro* hippocampus preparation [15]. These intrinsic theta rhythms appeared spontaneously without any pharmacological manipulations or artificial stimulation paradigms, and persisted even after the neighboring CA3 subfield was removed. It is thus clear that intrinsic theta frequency rhythms can be produced by local interactions between interneurons and pyramidal cells in the hippocampus. That is, the CA1 region of the hippocampus contains sufficient circuitry to be able to generate theta oscillations. Given this, using this *in vitro* hippocampus preparation presents an opportunity to understand cellular contributions to LFP theta. That is, we can remove several complicating factors by not needing to consider various pathways that exist in *in vivo* scenarios. Ambiguities can be greatly reduced and our ability to understand cellular contributions to LFP recordings is greatly enhanced.

Oriens-lacunosum/moleculare (OLM) cells are a major class of GABAergic interneurons with cell bodies in the outermost layer of the hippocampus (stratum oriens - SO) and perpendicular axonal projections to the innermost layer (stratum lacunosum-moleculare - SLM) inhibiting the distal apical dendrites of pyramidal (PYR) cells [16]. OLM cells play an important role in gating information flow in the hippocampus by facilitating intrahippocampal transmission from CA3 while reducing the influence of entorhinal cortical inputs [17]. Since OLM cells project to the distal dendrites of PYR cells they would be expected to generate large LFP deflections due to larger dipole moments [18]. However, these expectations may need to be modified since in addition to inhibiting distal dendrites in SLM, they can have an effect in stratum radiatum (SR), an inner to middle layer, since they inhibit interneurons that target PYR cells in SR [17].

In this paper we use computational modeling to determine the contribution of OLM cells to ongoing *intrinsic* LFP theta rhythms considering their interactions with local targets using the *in vitro* whole hippocampus preparation context. We take advantage of our previous modeling framework of inhibitory networks [19] to generate biophysical LFP computational models, and investigate the factors that influence theta LFP characteristics. By directly constraining our LFP models with experiment, we are able to predict the required connectivity profile between OLM cells and other inhibitory cells types, as well to show that OLM cells control the robustness, but not the power, of intrinsic LFP theta rhythms. We are also able to assess the spatial reach of the extracellular signal and so estimate the number of cells that contribute to the LFP signal. In general, we show how the many complex interactions lead to emergent LFP output that are non-intuitive and would not be possible to understand without biophysical LFP modeling in an experimentally constrained microcircuit context. As such, our work shows a way forward to obtain an understanding of cellular contributions to brain rhythms.

## Results

### Intrinsic theta rhythms in the hippocampus

A whole hippocampus *in vitro* preparation has been developed and shown to robustly and spontaneously generate *intrinsic* theta (3-12 Hz) rhythms [15]. An example of this intrinsic LFP rhythm is shown at the bottom of Fig 1. Considering this preparation, we previously estimated that a tissue size (i.e., network circuitry) of about one cubic millimetre is needed for the intrinsic theta rhythm to occur [20]. While it is clear that these intrinsic theta rhythms do not fully encompass *in vivo* theta rhythms, they undoubtedly exist without any special manipulations, and so are arguably part of the underlying biological machinery generating theta rhythms in the hippocampus. More importantly, to have a chance to understand the many different cellular contributions to LFP recordings, this preparation can be used to decipher the many interacting components.

**Fig 1.**
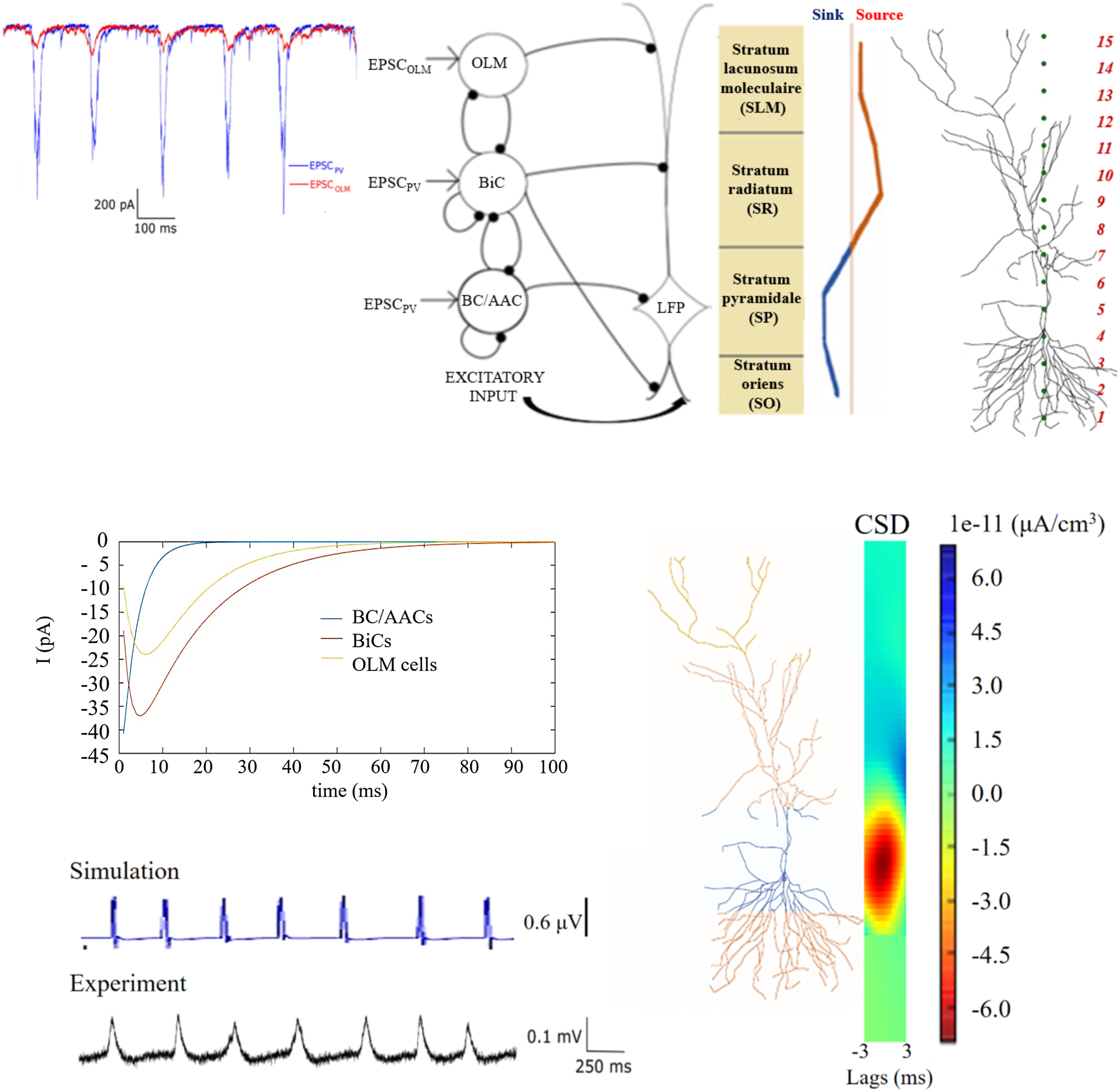
Model Setup and Experimental Essence. **Top:** A schematic of the network model used by [19] is shown in the middle. The network model contains single compartment representations for OLM cells, BiCs, and BC/AACs. Inhibitory synapses are represented by filled black circles. Each inhibitory cell receives excitatory post synaptic currents (EPSCs) that is taken from experimental intracellular recordings as shown on the far left (adapted from [19]). Each inhibitory cell synapses onto a PYR cell model as schematized. There are 350 OLM cells, 120 BiCs and 380 BC/AACs. Basal excitatory input is also included. An illustration of the polarity changes (source/sink) seen in the different labeled layers from LFP experimental recordings is shown on the right, and the detailed PYR cell morphology that is used along with the 15 equidistant electrode locations in the different layers is shown as red numbers on the far right. **Bottom:** IPSCs from the different cell types (colored as indicated) are shown on the left to show their different kinetics. Parameter values are given in Table 1, and the same coloring is used on the detailed PYR cell morphology to indicate the synaptic location regions for the different cell types. An example simulation of a computed LFP from the SR layer (using parameter values of g_sb_=6.00 nS and g_bs_=1.25 nS) is shown below, and the computed current source density (CSD) is shown on the right (averaged over time). On the very bottom is an example of an LFP recording from the SR layer (adapted from [19]).

**Table 1.**
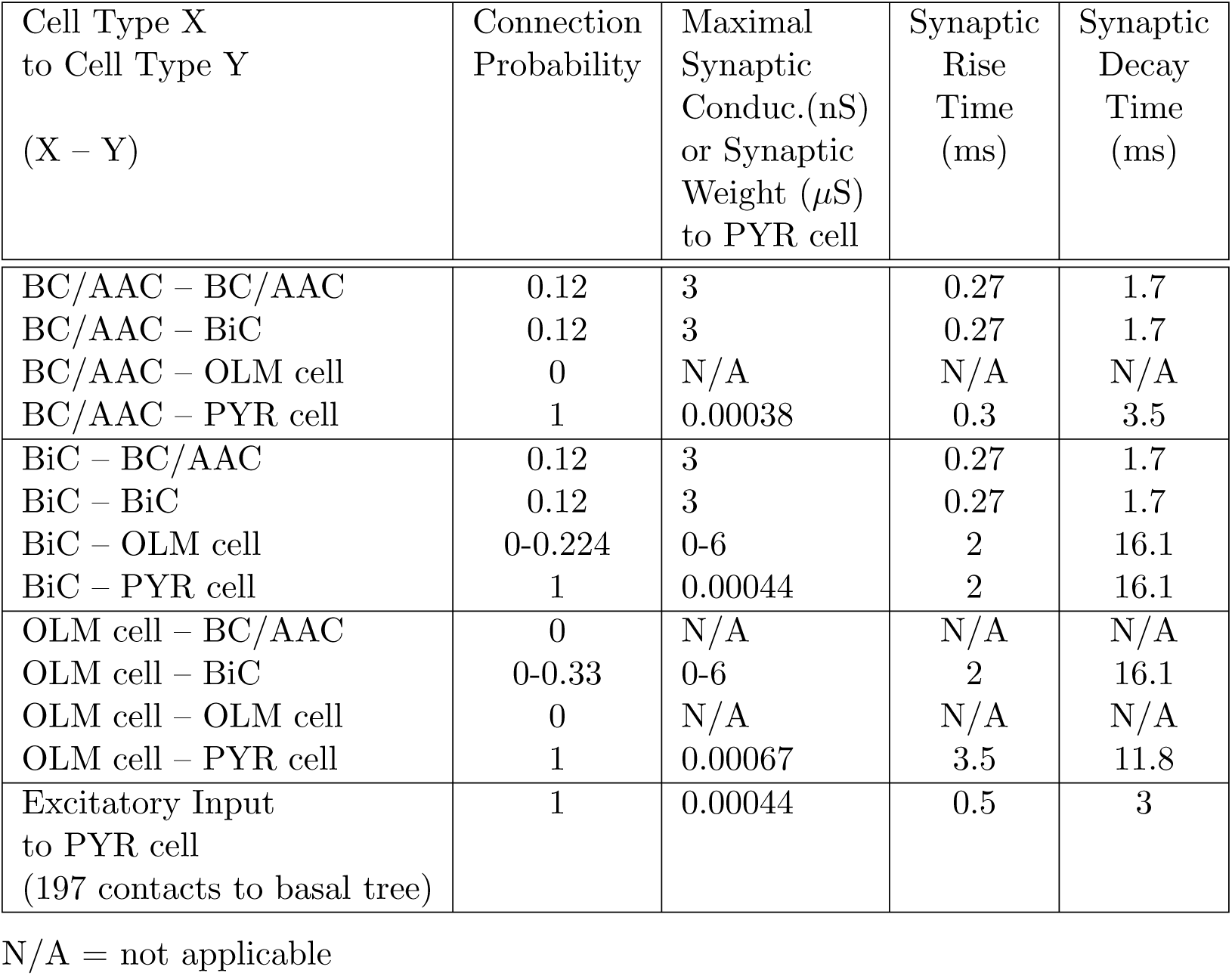
Connectivity Parameter Values.

| Cell Type X<br>to Cell Type Y<br>(X – Y) | Connection<br>Probability | Maximal<br>Synaptic<br>Conduc.(nS)<br>or Synaptic<br>Weight (μS)<br>to PYR cell | Synaptic<br>Rise<br>Time<br>(ms) | Synaptic<br>Decay<br>Time<br>(ms) |
| --- | --- | --- | --- | --- |
| BC/AAC – BC/AAC | 0.12 | 3 | 0.27 | 1.7 |
| BC/AAC – BiC | 0.12 | 3 | 0.27 | 1.7 |
| BC/AAC – OLM cell | 0 | N/A | N/A | N/A |
| BC/AAC – PYR cell | 1 | 0.00038 | 0.3 | 3.5 |
| BiC – BC/AAC | 0.12 | 3 | 0.27 | 1.7 |
| BiC – BiC | 0.12 | 3 | 0.27 | 1.7 |
| BiC – OLM cell | 0-0.224 | 0-6 | 2 | 16.1 |
| BiC – PYR cell | 1 | 0.00044 | 2 | 16.1 |
| OLM cell – BC/AAC | 0 | N/A | N/A | N/A |
| OLM cell – BiC | 0-0.33 | 0-6 | 2 | 16.1 |
| OLM cell – OLM cell | 0 | N/A | N/A | N/A |
| OLM cell – PYR cell | 1 | 0.00067 | 3.5 | 11.8 |
| Excitatory Input<br>to PYR cell<br>(197 contacts to basal tree) | 1 | 0.00044 | 0.5 | 3 |

N/A = not applicable

To examine the role of specific hippocampal interneurons in these intrinsic theta rhythms, Amilhon and colleagues [21] optogenetically activated and silenced interneurons expressing parvalbumin (PV) or somatostatin (SOM). PV cell types exhibiting fast firing characteristics include basket cells (BCs), axo-axonic cells (AACs) and bistratified cells (BiCs) [22]. OLM cells are SOM-positive but it is not the case that interneurons that express SOM are necessarily OLM cells. However, reconstructions of SOM cells in these studies with intrinsic theta were done to confirm that they were OLM cells [23]. Amilhon et al. [21] found that optogenetic manipulation of SOM cells modestly influenced the intrinsic theta rhythms. In contrast, activation or silencing of PV cells strongly affected theta. These results thus demonstrate an important role for PV cells but not SOM cells for the emergence and presence of intrinsic hippocampal theta, as given by the observed LFP recordings exhibiting theta rhythms.

LFP recordings in this preparation have a particular sink and source distribution in the different layers [15]. It is given by a single dipole characterized by positive deflections in stratum lacunosum/moleculare (SLM) and stratum radiatum (SR) and negative deflections in stratum pyramidale (SP) and stratum oriens (SO). The dipole is illustrated in Fig 1 (top right). This LFP laminar polarity profile is consistent across preparations. We note that in the whole hippocampus preparation the theta rhythms persist even when the CA3 region is removed, indicating that excitatory collaterals from CA3 are not a necessity for the emergence of the rhythm and the sink/source density profile. Thus, in our LFP model in this work, we assume that excitatory input to CA1 pyramidal (PYR) cells is restricted to the basal dendrites due to CA1 PYR cell collaterals [15].

### Previous network model framework as a basis

To try to understand how the complex interactions between different inhibitory cell types contribute to theta LFP rhythms, we had previously developed a computational network framework representing CA1 microcircuitry [19]. Driven by the ambiguous role of OLM cells in theta rhythms and the newly discovered connections between OLM cells and BiCs [17], we had developed network models to explore how OLM-BiC interactions influence the characteristics of theta rhythms. We took advantage of our previously developed PV fast-firing cell models [20], and developed OLM cell models [19] based on recordings from the whole hippocampus preparation. Because of distal contacts of OLM cells with PYR cells, we had used a multi-compartment PYR cell model to be able to incorporate this aspect in exploring the various interactions. The network model framework is shown in Fig 1 (top left) and a summary of the network model is provided in the Methods. We note that the network model was designed to explore cellular interactions and contributions to the ongoing intrinsic theta rhythms, and not to the generation of the theta rhythms per se. As such, all inhibitory neurons were driven by theta frequency inputs.

As schematized in Fig 1, the inhibitory cell populations encompass BC/AACs, BiCs and OLM cells that are driven by experimentally-derived excitatory postsynaptic currents (EPSCs). These EPSCs are from the ongoing rhythm and are of theta frequency (see Fig 1, top far left). Spiking output from the inhibitory cell populations lead to inhibitory postsynaptic currents (IPSCs) on the PYR cell. They are distributed on the PYR cell according to where the particular cell population targets. Thus, BC/AACs to somatic regions, BiCs to middle apical and basal regions and OLM cells to distal apical regions. IPSCs generated by the different cell types are shown in Fig 1 (bottom, left) (see Methods for details). In our previous work we used the spatial integration of the inhibitory postsynaptic potentials at the soma of a passive PYR cell model as a simplistic LFP representation [19]. This representation is in fact indicative of the intracellular somatic potential rather than the extracellular one, but it does allow the distal OLM cell inputs relative to more proximal PV cell inputs to be taken into consideration.

Using this computational model framework, we performed multiple simulations and showed that there are parameter balances that result in high or low theta power, and where OLM cells do or do not affect the theta power [19]. That is, OLM cells could play a small or large role in the resulting theta power depending on whether compensatory effects with BiCs occurred as a result of the size and amount of synaptic interactions between these cell types. Thus, interactions between OLM cells and BiCs in the CA1 microcircuitry seem to be an important aspect for the presence of intrinsic LFP theta rhythms. However, since we used an ad-hoc LFP representation, we could not do any direct comparisons with the experimentally recorded LFPs to decipher their output. Our ability to parse out the contribution of the different cell types or identify particular interactions was limited. Thus, while we were able to show that interactions between OLM cells and BiCs could play an essential role in the resulting theta power, we could not predict any particular parameter balances for this or extract possible explanations.

In the work here, we build on this model framework and develop biophysical LFP models. We use the inhibitory spiking output generated in [19] as a basis for generating biophysical LFPs, and we use the same PYR cell model. However, we now use the framework of volume conductor theory (see Methods) and generate actual extracellular potential output as a result of the overall activity of the inhibitory cell firings across the various layers of CA1 hippocampus. In addition we include excitatory input onto the basal dendrites to represent recurrent CA1 inputs (see schematic in Fig 1 and Methods for details) and directly compare with characteristics of experimental LFP recordings.

### Overall characteristics of biophysical LFP models

From our previous modeling study [19] we have several sets of inhibitory spiking output due to particular connection probabilities and particular synaptic conductances between OLM cells and BiCs. The connection probability from OLM cells to BiCs (*c_sb_*) varies from 0.01 to 0.33 with a step size of 0.02 producing 16 sets of connection probabilities; synaptic conductance values range from 0-6 nS for OLM cells to BiCs (*g_sb_*) and for BiCs to OLM cells (*g_bs_*) with a step size of 0.25 nS. Thus, for a given connection probability, there are 625 sets of spiking outputs from inhibitory cells, where each set represents a 850-cell inhibitory network with particular synaptic conductances. We consider a set to be a connectivity map representing the inhibitory cell populations.

For each connectivity map, a biophysical, extracellular LFP is generated. A virtual electrode probe is placed along the vertical axis of the PYR cell model to record its LFP output in a layer dependent manner. This PYR cell model is the “processor” of the LFP signal as it integrates postsynaptic inputs from different presynaptic populations. We compute LFPs at 15 equidistant sites along a linear axis - see Fig 1 (top far right). The PYR cell output corresponds to readouts of the postsynaptic activity elicited by the afferent interneuron, inhibitory cell populations that target the PYR cell in appropriate regions, referred to as the LFP “generator”. We note that although there is a single connectivity map representing the randomly connected inhibitory cell population, we perform several trials when randomly targeting the PYR cell to ensure the robustness of our results (see Methods for further details). To achieve effective electroneutrality, the extracellular sink needs to be balanced by an extracellular source, that is, an opposing ionic flux from the intracellular to the extracellular space, along the neuron; this flux is termed the ‘return current’.

We develop some initial intuition regarding the generation of our biophysical LFPs by computing them *without* including basal excitation. That way, all of the inputs received by the PYR cell model are inhibitory. Fig 2 illustrates the process and shows some examples. Let us first focus on the top left of Fig 2A. Next to each cell population in the network schematic are two examples of 1-second raster plots of spiking outputs (from the previously computed 5-second inhibitory network simulations in [19]) produced for particular parameter sets. These spikes give rise to IPSCs on the PYR cell model and the computed extracellular LFP at the somatic layer is shown next to the schematic. As shown, these particular parameter sets produce an LFP with positive deflections or with negative deflections. Let us next focus on the left of Fig 2B. One example of a 1-second raster plot is shown, and for this parameter set, the LFP has only a few positive deflections. Assuming that one population burst in the raster plot leads to a single peak in the LFP, there would be 29 peaks in the LFP for a 5-second simulation (i.e., about 5.8 Hz frequency). Note that the raster plots for this example are not very different from the examples shown in Fig 2A. We compute LFPs at all layers as represented by the 15 virtual electrodes shown in Fig 1 (top far right) for the 625 sets of inhibitory spiking outputs across *g_sb_* and *g_bs_* values at a particular connection probability *c_sb_*. The colored plot at the top right of Fig 2A shows the polarity of the LFPs at the somatic layer, and the color plot in Fig 2B shows the number of LFP peaks in the somatic layer. In Fig 2C normalized spike numbers for all interneuron populations are shown.

**Fig 2.**
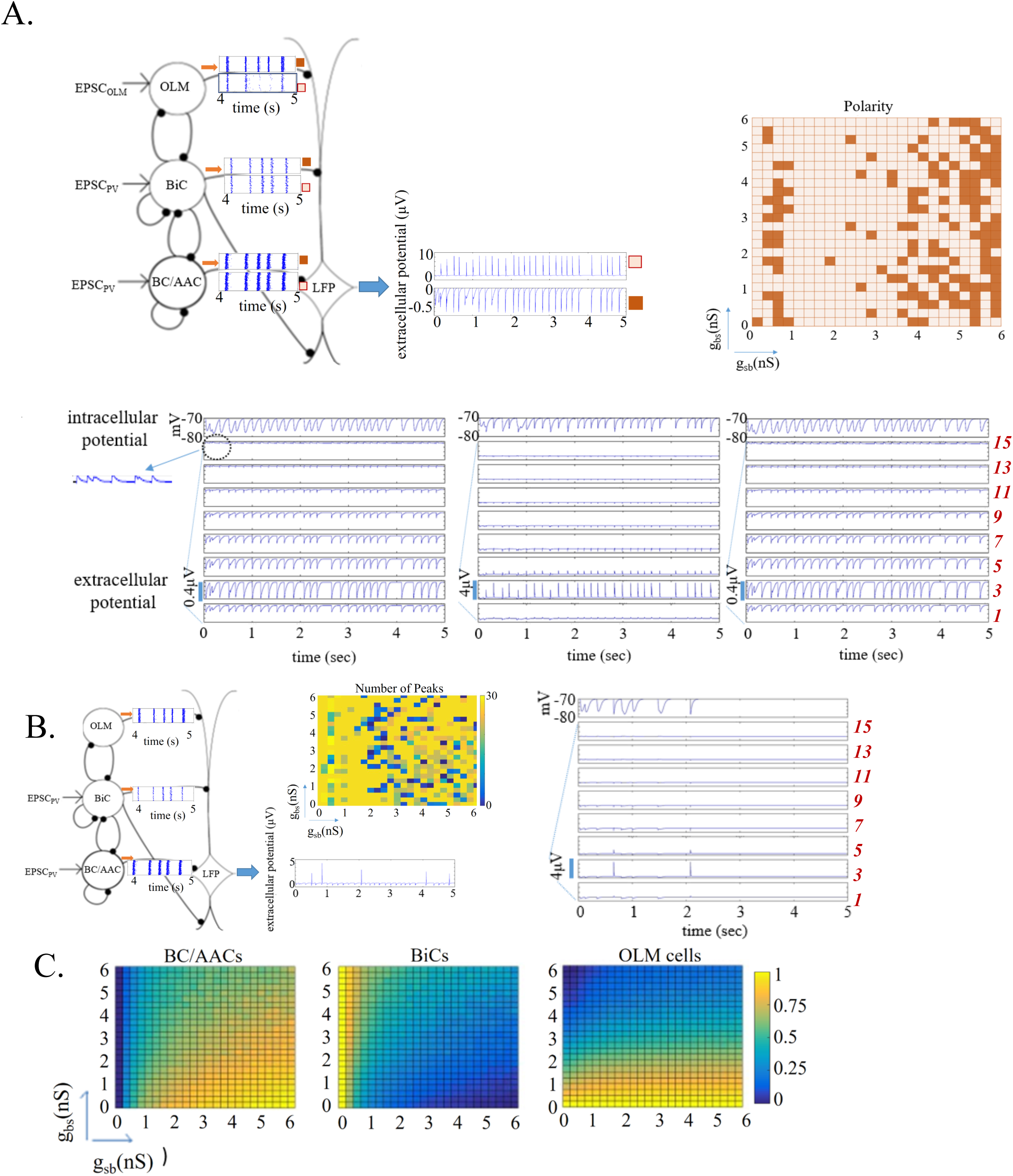
Biophysical LFP Computation: Features and Examples. LFPs are shown in the absence of basal excitatory input and for c_sb_=0.21. **A. Top:** Schematic includes 2 raster plot example outputs for the given inhibitory cell population, and the resulting LFPs at the somatic layer, with positive and negative deflections for these 2 examples. Parameter values are g_sb_=1.5, g_bs_=-5.5 nS for positive and g_sb_=0.5, g_bs_=0.75 nS for negative deflections. The color plot on the right shows the polarity at the somatic layer, SP, electrode 4. Negative polarity:dark-colored, positive polarity:light-colored. **Bottom:** LFP output for all layers are shown for 3 examples where the polarity is negative, positive and negative at electrode 3 (left to right). Parameter values are (left to right): g_sb_=0.5, g_bs_=0.75; g_sb_=1.5, g_bs_=5.5; g_sb_=5.75, g_bs_=0.75 nS. Inset shows a blow up of LFP output at electrode 13 (SLM) to show positive deflections. Also shown is the intracellular somatic potential of the PYR cell. **B.** Schematic includes 1 raster plot example, and the resulting LFP output at SP has 5 peaks. A maximum of 29 peaks is possible (see text). Parameter values are g_sb_=2, g_bs_=0.75 nS. The color plot shows the number of peaks that appear in the 5-second LFP computation at SP, electrode 4. On the right is an example of LFP output for all layers as well as the intracellular somatic output which also shows a loss of peaks. Parameter values are g_sb_=2.25, g_bs_=5.0 nS. **C.** Interneuron activity for each interneuron population, normalized such that the number of spikes for a given pair of synaptic conductances is divided by the maximal number considering all pairs of synaptic conductances. Maximal number (5-second trace): 16, 327 (BC/AACs), 6, 808 (OLM cells), 4, 589 (BiCs).

As a first approximation, given the network model framework and previous work we can say the following about the LFPs: Those governed mainly by synaptic inputs and not return currents are characterized by narrow waveform shapes as the synaptic inputs from any particular interneuron population enters the PYR cell in a synchronized fashion. This is due to the inhibitory cells in a given population being driven by rhythmic EPSCs that give rise to coherently firing inhibitory cells in a given population (see example raster plots). We note that the EPSCs that were used in the simulations are not perfectly synchronized since the measured experimental variability was included in designing the EPSC inputs to use in the inhibitory network simulations (see Methods). On the other hand, return currents constitute a summation of less synchronized exiting currents that originally entered the cell at different locations. Therefore, LFP deflections governed by return currents are generally wider occurring also with certain latencies. Further, we would expect that the LFP recorded from different layers would first and foremost be influenced by the interneurons that project to that region. We also note that the width of the LFP deflection would not only be influenced by the nature of the current (synaptic inputs or return currents) but also by the synaptic time constants defining the shape of the IPSCs. IPSCs for the different cell populations are shown in the middle of Fig 1 where it can be seen that the IPSCs produced by OLM cells and BiCs are wider relative to the IPSCs from BC/AAcs. Thus, we expect that positive LFP deflections would be recorded in locations where OLM cells, BiCs and BCs project, with wider LFPs for OLM cell projection locations, and that LFPs dominated by return currents would be recorded in locations where there are no direct inputs from interneurons. However, due to interactions between BiCs and OLM cells, this is not necessarily the case as return currents from distant interneuronal inputs may prevail in regions where other interneurons directly project. In fact, interactions between OLM cells and BiCs can strongly modulate the relative balance between synaptic inputs and return currents, which in turn can strongly modulate the distribution of sinks and sources in the resulting LFP.

The two examples of LFP output at the somatic layer in Fig 2A show one with narrow positive deflections and the other with wider negative deflections. This thus indicates that the BC/AAC inputs that synapse at the somatic layer dominate for the positive deflection LFP example whereas BiC and OLM cell inputs that synapse more distally dominate for the negative wider deflection LFP example. The example in Fig 2B of LFP output at the somatic layer indicates that a loss of peaks can occur due to the superposition of synaptic inputs and return currents. Another “loss of peaks” example is shown on the right of Fig 2B, and LFP output from multiple layers is shown in addition to the intracellular somatic output. For this example, the peak loss is also partially reflected in the intracellular somatic output. However, loss of peaks in the LFP output is not necessarily reflected in the somatic intracellular recording. Note that since the PYR cell is only receiving inhibitory input in these set of simulations, somatic intracellular potentials always have negative deflections. How the extracellular potential features change as a function of the synaptic conductances between BiCs and OLM cells is summarized in the color plots of Fig 2A for the polarity and Fig 2B for the number of peaks (somatic layer).

Let us consider the colored plot of Fig 2A. We find that we can approximately distinguish four regions as the *g_sb_* conductance is increased. For small *g_sb_* values (0-1 nS) the amount of inhibition that the BiCs receive from the OLM cells is minimized allowing the BiCs to be at the peak of their activity (see Fig 2C). Consequently, the inhibition that the OLM cells and BC/AACs receive from the BiCs is maximized causing their activities to be minimized (see Fig 2C). As a result, the extracellular potential in the somatic region is governed by return currents leading to negative polarity LFPs in the somatic layer (i.e., mainly dark-colored in Fig 2A plot), primarily due to the BiC synaptic inputs on the ‘middle’ region (SR layer) and ‘basal’ region (SO layer) of the PYR cell. As we increase *g_sb_* (1-2.25 nS), we encounter mainly positive polarity LFPs (ie., light-colored). In this region the inhibition onto the BiCs is increased and thus their activity is decreased, as can be seen in Fig 2C, causing a decrease in the amount of the inhibitory current onto the PYR cell from BiCs. As a result, the magnitude of the return currents caused by the BiC synaptic inputs is decreased at the somatic layer. Simultaneously their ability to inhibit the BC/AACs is also decreased so that the BC/AACs become more active and their direct inhibition onto the PYR cell also increases. Since both BiCs and OLM cells activity is low in this region while BC/AAC activity is increased, the somatic LFP is governed by BC/AAC inputs rendering the extracellular LFP positive. As we further increase *g_sb_* (2.25-5 nS) the silencing of the BiCs increases even further and their ability to silence the BC/AACs is further reduced. Simultaneously OLM cell activity increases. Thus the somatic LFP is influenced by direct synaptic inputs from BC/AACs and also return currents from OLM cells (sparse dark-coloring in this region). Interestingly, the majority of the “loss of peaks” in somatic LFP output occurs in this region (see blue-green regions in the color plot in Fig 2B) since the superposition of synaptic inputs and return currents occurs most often in this region. That is, cancellations occur even leading to abolishment of the entire rhythm. Finally, for *g_sb_* from 5.25-6 nS, the BiCs are maximally inhibited and BC/AACs are at the peak of their activity. While we might expect domination from the BC/AAC synaptic inputs for these values, it turns out that return currents (negative polarity) dominate. This can be explained by the increased activity of OLM cells which are also at the peak of their activity producing strong return currents in the somatic region. In summary, light-colored regions signify that BC/AACs dominate the extracellular somatic potential and dark-colored regions signify that other inhibitory cell types (BiCs or OLM cells, or both) contribute more strongly.

At the bottom of Fig 2A, we show three examples of LFP recordings at multiple layers as well as the somatic intracellular potential, for increasing values of *g_sb_* from left to right. To allow an appreciation of the changing magnitude of the signal, we use the same resolution on the ordinate axis for all LFP plots shown. On the left (*g_sb_*=0.5 nS) we see that the signal is governed by return currents (negative polarity) in the entire SP (electrodes 3 and 5), in SO (electrode 1) and in SR (electrodes 7,9 and 11). Synaptic events govern SLM (electrodes 13 and 15) where OLM cells directly project leading to positive polarity. In the middle (*g_sb_*=1.5 nS), the LFP in SP and SO is governed by synaptic inputs (positive polarity), and in SR and SLM by return currents (negative polarity). As expected, we find that the positive polarity LFP in SP here is narrower relative to the positive polarity LFP in SLM on the left, because the IPSCs produced by OLM cells are wider relative to those of BC/AACs, as shown in Fig 1 (middle). On the right where *g_sb_*=5.75 nS, we observe a similar trend as for the example on the left where *g_sb_*=0.5 nS with return currents dominating.

We would like to use our computational LFPs to determine how the different inhibitory cell types contribute to theta LFPs as recorded experimentally in the *in vitro* whole hippocampus preparation. As described above, our overall network model (Fig 1) is intended to capture an intrinsic theta rhythm in the CA1 region of the *in vitro* preparation. CA3 input is not required but local excitatory input which occurs on basal dendrites [24] does need to be included. To include this, we take advantage of previous modeling studies [19, 25] as detailed in the Methods. Including excitatory input would clearly affect resulting biophysical LFP outputs. We expect that the LFP amplitude in SO might decrease even further in the presence of basal excitation as excitatory and inhibitory BiC inputs could cause mutual cancellations in this region. As return currents mostly exit close to the somatic region where the surface area is larger, the effect of basal excitation is expected to be stronger in SO and SP since most of the current would be expected to have exited before reaching SR and SLM. In general, we expect there to be a range of possible LFP characteristics based on the above LFP computations done in the absence of basal excitation. We expect that the addition of excitatory input will influence the LFP in non-intuitive and nonlinear ways and the intuition developed above will be helpful in deciphering the contribution of the different cell populations to the LFP.

### Prediction of synaptic conductances and connection probabilities between BiCs and OLM cells

Although we would like to understand how different cell populations contribute to intrinsic theta LFP rhythms in general, we focus on OLM cells in this work. Our model network framework was developed based on knowing that connections exist between BiCs and OLM cells [17]. Given this, there are two pathways to consider for how OLM cells could influence ongoing intrinsic theta LFP rhythms. They can influence LFP output indirectly through disinhibition of proximal/middle dendrites of the PYR cell (OLM-BiC-PYR pathway), and this influence would act on top of the direct inhibition that OLM cells exert on distal, apical dendrites (OLM-PYR pathway). As shown above, many different LFP features can be exhibited in the absence of basal excitation (see Fig 2). It is interesting to note that our biophysical LFP output does not necessarily exhibit theta frequencies, despite being driven by theta frequency EPSC inputs (see Fig 2B). This underscores the importance of modeling biophysical LFPs as the interaction of synaptic and return currents on the extracellular signal can strongly affect the resulting LFP frequency.

We now proceed to include basal excitation and perform a full set of computations for all connection probabilities (*c_sb_*) and synaptic conductances (*g_sb_*, *g_bs_*). With these computed biophysical LFPs in hand, we do direct comparisons with experimental LFPs from the whole hippocampus preparation *in vitro*. Specifically, we classify each set of network parameters as *selected* or *rejected* based on whether our computed LFPs are able to reproduce two robust characteristics exhibited experimentally. These are: (i) the laminar polarity profile exhibits a single dipole with sinks in the basal dendrites and sources in the apical dendrites, and (ii) the frequency of the LFP traces across all layers is in the theta frequency range. These characteristics are shown in Fig 1. We note that our model setup in which experimentally-derived theta frequency EPSCs are input to the inhibitory cells means that the LFP rhythm should have a theta frequency. However, as we have shown above, the resulting biophysical LFP frequency can be much less due to synaptic and return current interactions and cancellations (see Fig 2B). Specifically, the frequency of the EPSCs used from experiment is about 5.8 Hz. Thus, in enforcing the theta frequency on our LFP computations, it is only necessary to impose a lower bound. We use 3 Hz to be similar to experiment [15]. In Fig 3 (top) we show an example of computed LFPs across the different layers for a parameter set that was selected. The bottom of Fig 3 shows LFP outputs for three different parameter sets that were rejected - incorrect polarities and frequencies are apparent. Note that ordinate resolutions are adjusted across the layers so that the frequency and polarity of computed LFPs can be readily seen in each layer in doing the comparison.

**Fig 3.**
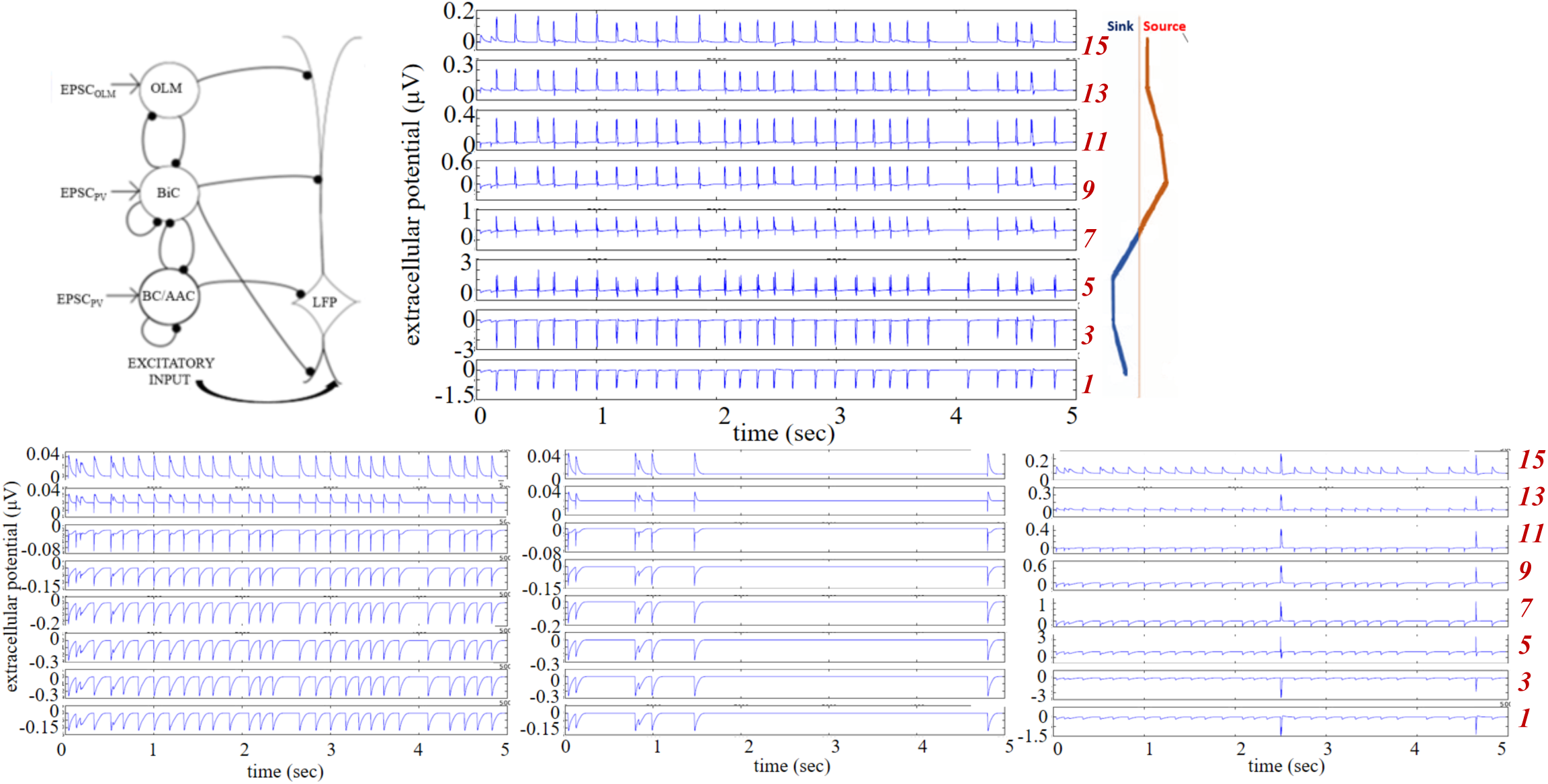
Example LFPs from Selected and Rejected Parameter Sets. Computed LFPs are shown across multiple layers. **Top:** Selected parameter set: g_sb_=6, g_bs_=1.25 nS. **Bottom:** Rejected parameter sets (left to right): g_sb_=0.5, g_bs_=0.75 nS; g_sb_=0.5, g_bs_=3.5 nS; g_sb_=2.5, g_bs_=1 nS. c_sb_=0.21 for all.

Classifying each parameter set, we summarize our results in Fig 4 where selected parameter sets are shown in purple and rejected ones in yellow. We observe the following: For low *c_sb_*, the plots have a checkered appearance since small changes in *g_sb_* and *g_bs_* cause the system to alternate between being selected or rejected. As *c_sb_* increases, there is a clearer separation in (*g_sb_*, *g_bs_*) parameter space of selection or rejection. This is observed from *c_sb_*=0.19 to *c_sb_*=0.25. In this range, we consider the system to be robust as it is not very sensitive to synaptic conductance perturbations. However, for *c_sb_*=0.19, 0.23 and 0.25, the selected parameter sets are quite narrow. As *c_sb_* is further increased, the checkered patterning returns. Note that the selected sets are mainly affected in one direction as *c_sb_* changes. That is, across *g_sb_* rather than *g_bs_* values. Further, we note that doing this classification, it is more the polarity rather than the frequency of the LFP signal that delineates selected and rejected parameter sets. This is shown in S1 Fig where we do not apply any frequency bound or use different lower frequency bounds. While there is some change in selected and rejected parameter sets, they are minimal.

**Fig 4.**
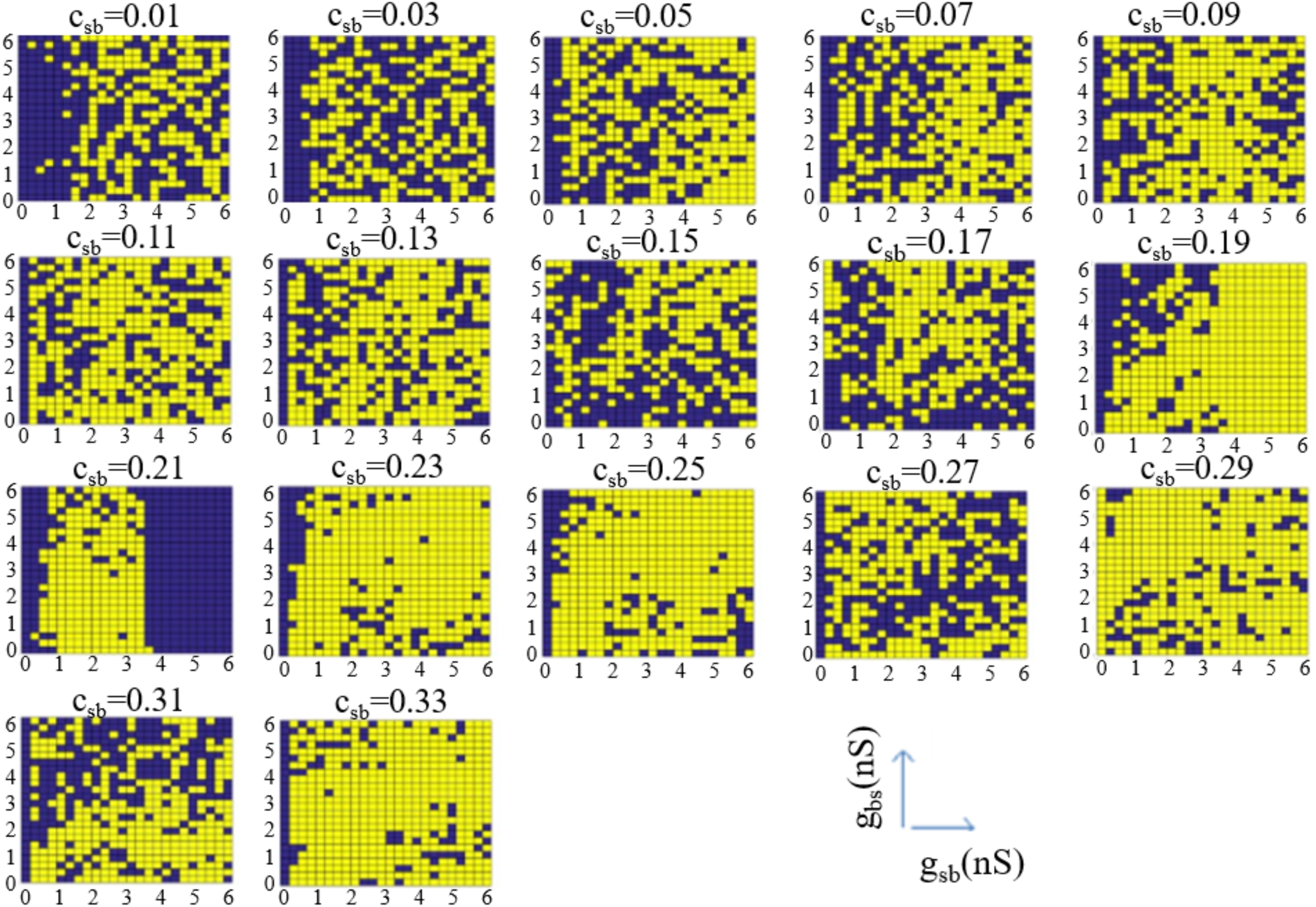
All Selected and Rejected Parameter Sets. Parameter sets are considered as selected (purple) if computed LFPs match LFPs from experiment in polarity and frequency (3 Hz lower bound). Otherwise, as rejected (yellow). A clear separation in parameter space occurs for c_sb_ = 0.21.

Since there is natural variability in biological systems, we assume that sensitivity to small perturbations in parameter values is anathema to having robust LFP theta rhythms. Noting that the synaptic conductance resolution in our simulations is 0.25 nS, and that a minimal synaptic weight can be estimated as larger than this (see Methods), we consider that (*g_sb_*, *g_bs_*) parameter sets that do not yield at least two complete, consecutive rows or columns of purple (selected) are inappropriate for biological systems. That is, variability that is less than a minimal synaptic weight would not make sense. Looking at this in Fig 4, we first note that there is never at least two complete purple rows for any *c_sb_*, but there are cases of two or more complete purple columns, namely, *c_sb_*=0.03 and 0.21. A complete purple column for *g_sb_*=0 is invalid since OLM to BiC connections exist [17]. Thus, *c_sb_*=0.03 can be eliminated leaving *c_sb_*=0.21 as appropriate. For this connection probability, the transition from selected to rejected networks and vice versa strongly depends on *g_sb_* rather than on *g_bs_* values, revealing a more important role for the former. In summary, by directly comparing characteristics of our computed biophysical LFPs with those from experiment, we predict a connectivity of *c_sb_*=0.21, *g_sb_* values of 3.5 to 6 nS, and the full set of *g_bs_* values (*g_sb_* ≠ 0, *g_bs_* ≠ 0). We will refer to this set of parameter values as the *predicted* regime. In Fig 5 we show example LFP responses across several layers for a set of parameter values from this *predicted* regime.

**Fig 5.**
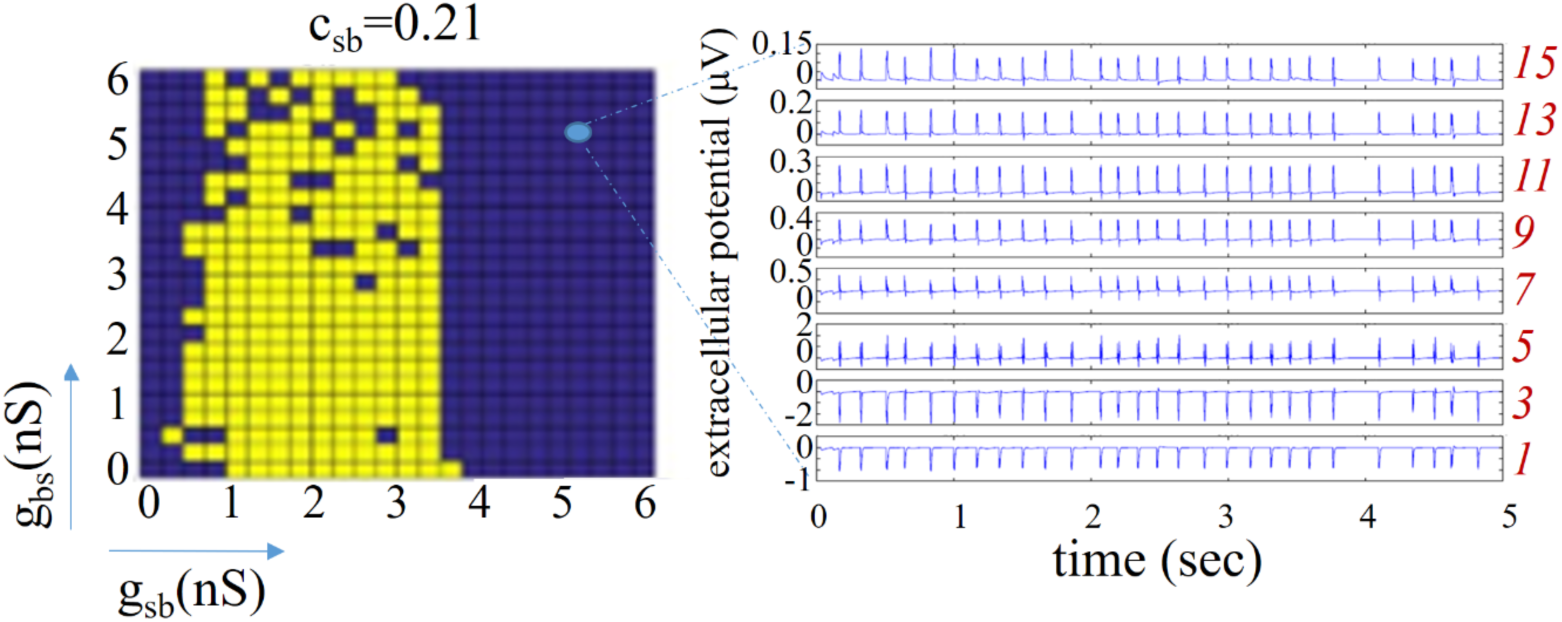
Predicted Regime. For c_sb_ = 0.21, selected parameter sets (purple) include g_sb_ values of 3.5-6 nS, and all g_bs_ values. Rejected sets are in purple. On the right are LFP traces from 8 electrodes for a parameter set of g_sb_ = 4.75, g_bs_ = 4.50 nS.

### OLM cells ensure a robust theta LFP signal, but minimally affect LFP power, and only through disinhibition

We continue our analysis but now we focus only on the *predicted* regime. We decompose the signal to be able to examine the contribution of the interneuron subtypes to the power of the LFP. We separate our interneuron subtypes into two groups - PV subtypes which are BC/AACs and BiCs, and OLM cells. These two groups are represented by distinct mathematical models of fast-firing (PV) and somatostatin-positive (SOM) inhibitory cells based on whole cell recordings from the whole hippocampus preparation [19]. We perform spectral analyses of our computed LFPs and use the peak amplitude (which is always within the theta range) as a measure of the power of the theta network activity. The peak power is computed for each of the 15 electrodes (i.e., all layers), and we plot the maximum value from all of the layers in the color plots of Fig 6. This is illustrated on the right of Fig 6A. We first simulate the spectral LFP power when all presynaptic inhibitory cell populations are present. As shown in Fig 6A, a robust power feature emerges. When all presynaptic origin populations are present, the predicted regime shown in purple in Fig 5, produces LFP responses whose power shows minimal variability. This is an interesting observation on its own, as the power of the LFP varies little across hippocampus preparations [15]. Thus, our predicted regime satisfies another characteristic of experimental LFPs. For completeness, we show peak power computations for all connectivities in S2 Fig.

**Fig 6.**
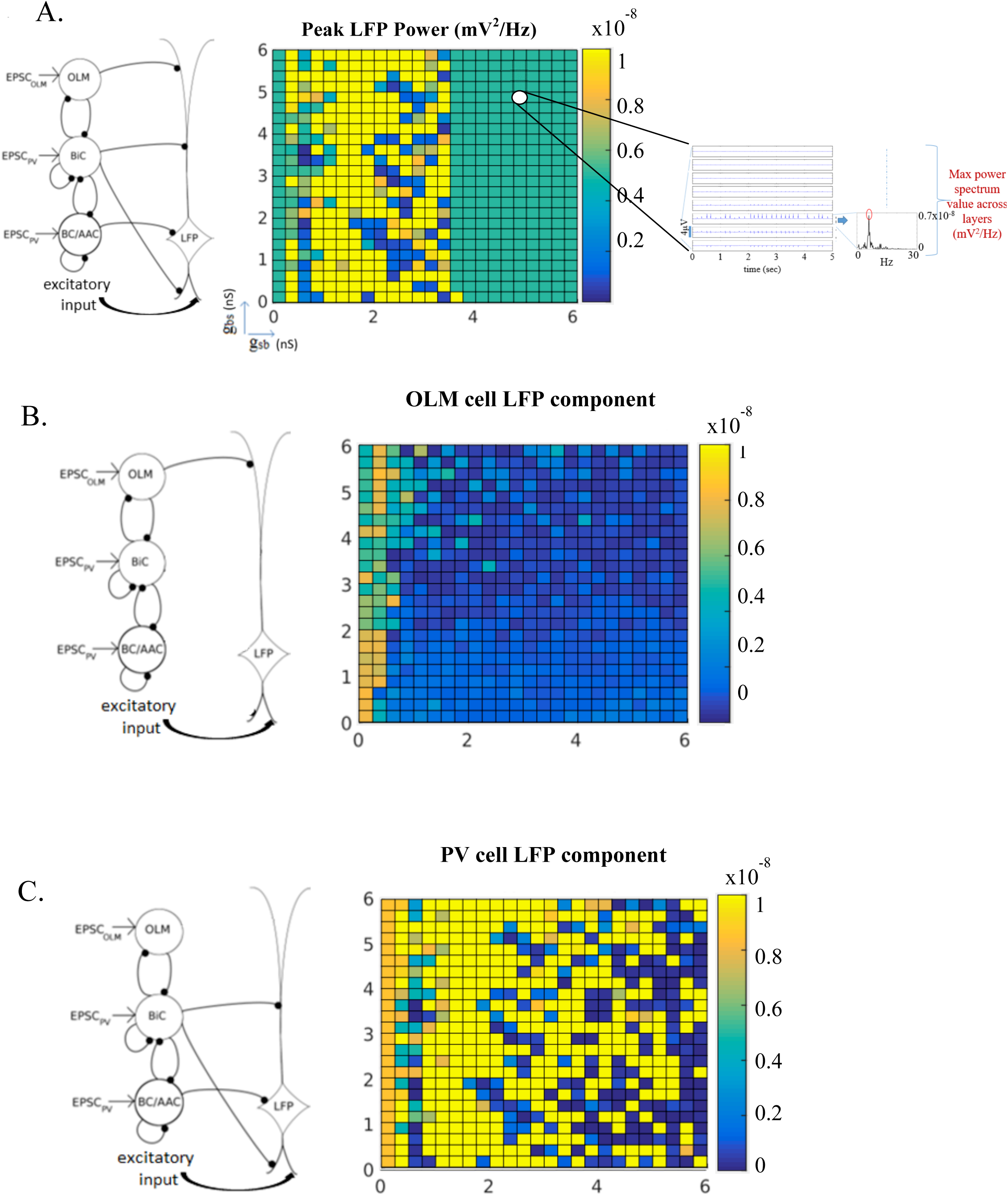
Decomposition of the LFP Signal. **A.** All presynaptic cell populations are present. **B.** Only OLM cells are present. **C.** Only BiCs and BC/AACs are present. Schematics on the left show the cell populations projecting to the PYR cell. Color plots are of peak power computations across g_sb_ and g_bs_ parameter values. See illustration to the right of A for this computation (see text for details).

To examine the role of presynaptic origin populations on the LFP we decompose the signal by selectively removing OLM to PYR cell connections or PV to PYR cell connections and then computing and plotting the peak power as described above. Selective removal of synapses from PV cells to the PYR cell yields an LFP response whose presynaptic origin population is due to the OLM cell population. The resulting LFP power is low and depends weakly on *g_bs_* (Fig 6B). This shows that OLM cells minimally contribute to the signal power as a presynaptic origin population. Viewing this from a broader perspective, these results indicate that disinhibition of non-distal apical dendrites via an OLM-BiC-PYR pathway plays a much larger role relative to a direct OLM-PYR pathway in producing the LFP power. Along the same lines, disinhibition of distal dendrites through a BiC-OLM-PYR pathway does not have much of an effect on LFP power. Fig 6C shows the result when we selectively remove the synapses from OLM cells to the PYR cell to yield an LFP response whose presynaptic origin is the PV cell population. It is clear from the magnitude of the signal power that it is indeed mainly due to the component from PV cells. Interestingly, the previously seen robustness when all presynaptic cell populations were present (Fig 6A) is now lost. To quantify all of this, we computed the mean and standard deviation (std) of the peak powers in the predicted regime for Fig 6A–C. Respectively, they are (mean, std): (5.1 × 10^−9^, 1.7 × 10^−23^), (9.7 × 10^−10^, 5.6 × 10^−10^), (2.6 × 10^−8^, 3.8 × 10^−8^). When all of the cell populations are present, there is minimal variability, and when the PV cell populations are removed, the average power decreases five-fold and there is some variability. However, when only PV cell populations are present, there is an increase in the average power and the variability is large. It seems clear that OLM cells do not contribute much to the average LFP power but removing their inputs prominently affects the robustness of the LFP signal. Therefore, we propose that OLM cells have the capacity to regulate robustness of LFP responses without affecting the average power.

In a recent study, Amilhon and colleagues [21] showed that SOM cells (putative OLM cells) do not appear to play a prominent role in the generation of intrinsic LFP theta rhythms since there was only a weak effect on LFP theta power when they optogenetically silenced SOM cells. Our results are in agreement with this observation. As shown in Fig 6B, the contribution of OLM cell inputs to the LFP power is small. However, if we want to make a more accurate comparison to Amilhon and colleagues’ OLM optogenetic silencing experiments we should compare the power of the LFP in the predicted regime in Fig 6A (mean value of 5.1 × 10^−9^) to the power of the LFP in Fig 6C for *g_sb_*=0 and *g_bs_*=0 when OLM-PYR connections are also removed (8.5 ×10^−9^). They are clearly comparable. It is interesting to note that it is already apparent from Fig 6A that OLM cells minimally affect LFP power. Consider that for the parameter regime of *g_sb_* = 0 and across all *g_bs_*’s, the LFP power magnitude is the same (5.1 ×10^−9^) as the average power of the predicted regime. In this *g_sb_* = 0 parameter regime, OLM cell to BiC connections are not present but the OLM cell to PYR cell connections are still present so that OLM cells can still contribute to the LFP response via a direct OLM-PYR cell pathway. Given that the power does not change indicates that any LFP power contribution due to OLM cells occurs mainly via the indirect OLM-BiC-PYR pathway. Overall, our results show that OLM cells do participate but in such a way that their presence would be unnoticed if one were only measuring LFP power.

To gain insight into how OLM cells affect the robustness of the LFP signal, we examine further what is revealed with our LFP decompositions. In Fig 7 we show the peak power plots for the PV cell (Fig 7A) and OLM cell (Fig 7B) decomposition components in which the non-predicted regime is overlaid with gray. For each component, we show three examples of LFP responses across the different layers, and raster plots that correspond to each cell population for the last second of the simulation. These examples illustrate the various cases of impaired LFP responses that occur when OLM or PV cell connections to the PYR cell are removed. It is also evident that the different LFP patterns cannot be intuited from the raster plots alone.

**Fig 7.**
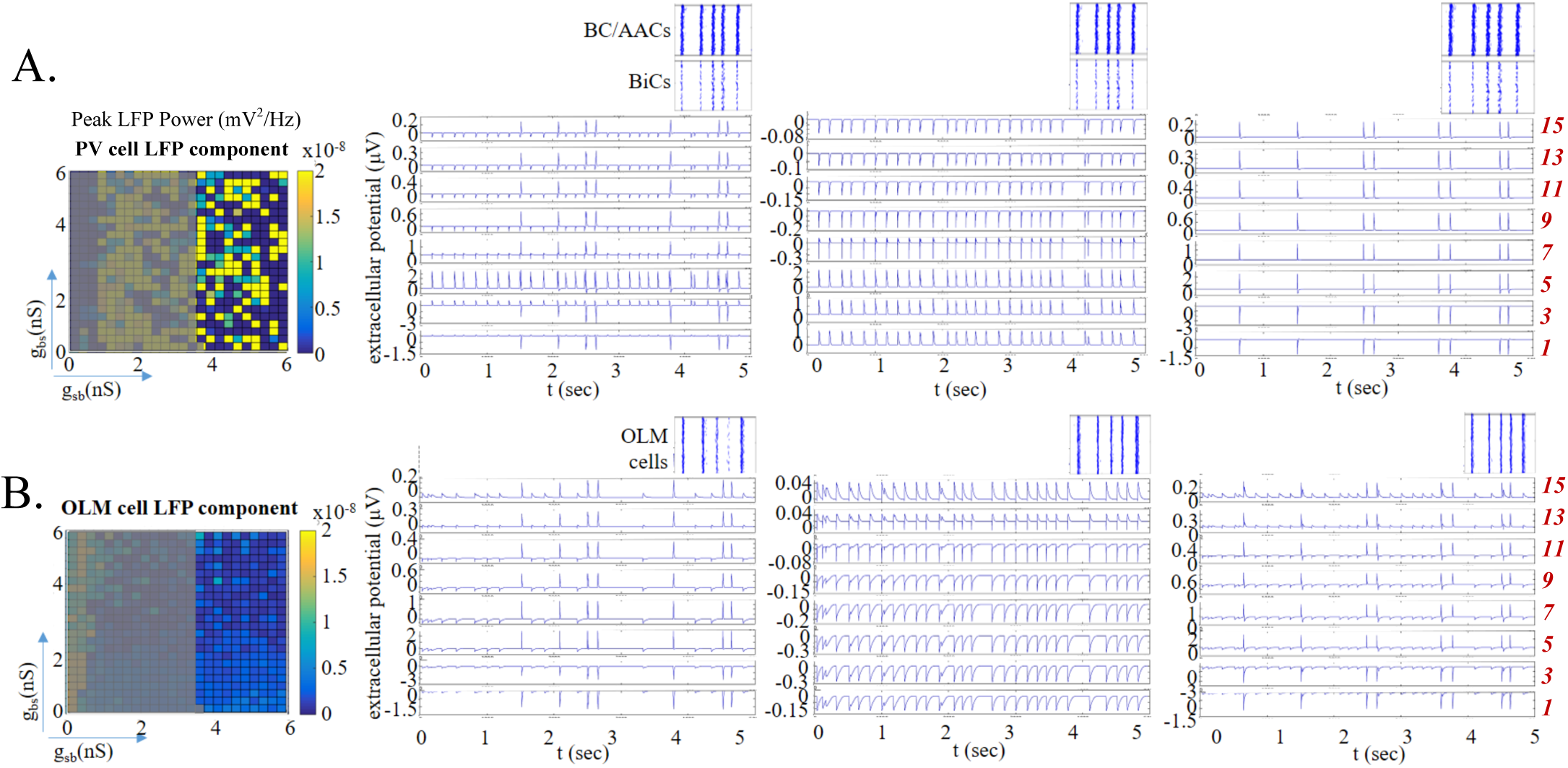
LFP Pattern Examples In Predicted Regime When Only Either PV or OLM Cell Populations Are Present. Peak power color plots as in are shown but with a different color resolution. A gray overlay is added to the plots to emphasize the predicted regime. Three examples of LFP responses (5 sec) across the different layers are shown to illustrate the different patterns observed. For each example, spike rasters (1 sec) for the particular example are shown for PV cells (BiCs and BC/AACs) or OLM cells. **A.** PV cell LFP component. **B.** OLM cell LFP component. Parameter values for left, middle and right columns are respectively: (g_sb_, g_bs_) = (5, 2.75), (5.5, 0.5), (5.75, 1) nS.

For the middle LFP response examples (low *g_bs_* and high *g_sb_*) of Fig 7, we note that OLM cells and BC/AACs have maximal activities and BiCs have minimal activities (see Fig 2C). Thus, synaptic current influences are obvious at the layers where OLM or BC/AACs contact, and return currents at other layers. Inappropriate polarity across the layers is manifest. This pattern of impaired LFP response occurs in about a quarter of the PV cell LFP component parameter sets, and in less than half of the OLM cell LFP component parameter sets. For the PV cell LFP component, most of the other parameter sets yielded LFP responses in which there was no rhythm, as shown in the right example of Fig 7A. Interestingly, in the rest of the cases (less than a third) there is a loss of rhythmicity in all layers except for the somatic layer as illustrated in the left example. These patterns show that there is an ongoing ‘battle’ between basal excitation and PV cell inputs that can yield a wide range of LFP powers from low (no rhythm - right example) to high (left and middle examples). For the majority of the OLM cell LFP component parameter sets, there is a loss of rhythmicity as shown in the left and right examples of Fig 7B. From the temporal profile and polarity, it is clear that the high amplitude LFP peaks are due to basal excitatory inputs. For larger *g_bs_* values OLM cells are less active (see Fig 2C) and LFP responses across the layers become dominated by peaks due to basal excitation rather than synaptic and return currents due to OLM cells. Overall, cancellations and rhythm loss occurs due to interactions between OLM cells’ synaptic and return currents and excitatory inputs. As summarized in the peak power plots of Fig 6C or Fig 7A, PV cell inputs alone are not capable of sustaining the robustness throughout the predicted regime and the impaired LFP signals show a large variability. With OLM cell inputs alone, there is low LFP power either because of loss of rhythmicity or because of low amplitude rhythms (Fig 6B or Fig 7B peak power plots).

### With and without basal excitation

As one might expect, including basal excitation to incoming inhibitory inputs from different cell populations adds to the complexity of untangling nonlinear, interacting components producing the LFP. We relied on our developed intuition when basal excitation was not included (Fig 2) and our LFP decompositions to help reveal the different roles that OLM cells and PV cells play in LFP theta rhythms. Thus, in finding that the LFP power is a robust feature in the predicted regime of synaptic conductance and connection probabilities, we are able to understand that it is critically the OLM cell population that brings about this robust feature. However, this robust feature is apparent only when basal excitation is included. This is clearly visualized in Fig 8 where we plot the peak power color plots with and without basal excitation when all cells are present and with OLM cell and PV cell LFP components. Removal of basal excitatory inputs in the case where all cells where present (Fig 8, top) leads to a loss of robustness. The mean and std in the predicted regime without basal excitation is 6.2 × 10^−9^ and 8.0 × 10^−9^ respectively. While the mean is comparable to when basal excitation is present, the standard deviation is much larger (see values with basal excitation above). Co-activation of inhibition and excitation is clearly important for this robust feature to emerge.

**Fig 8.**
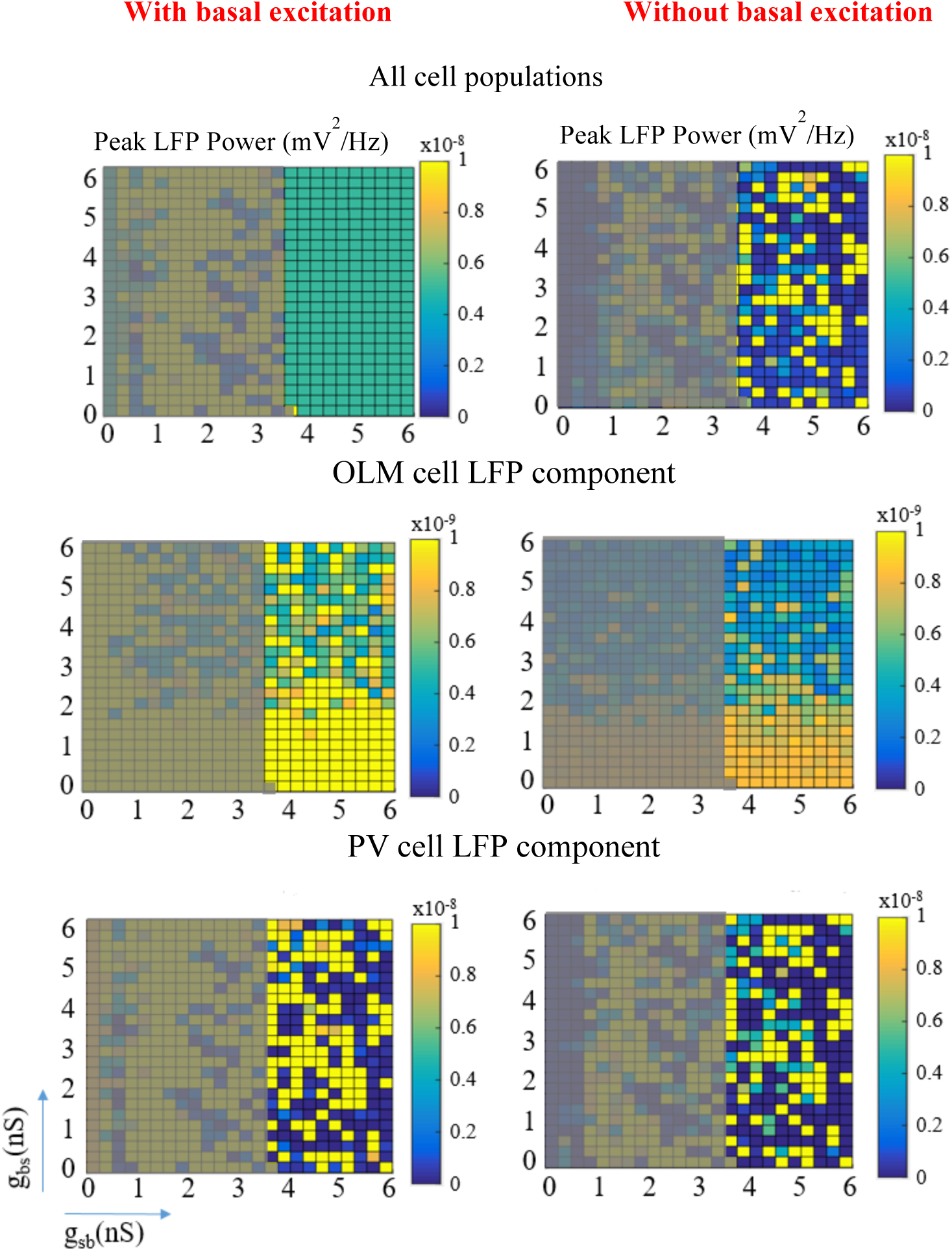
Peak Power Plots With and Without Basal Excitation. Peak Power Plots With and Without Basal Excitation. The color plots represent peak power as described in Fig 6 and with a gray overlay as in Fig 7. Note that different color resolutions are used here to facilitate comparison for particular cell populations (i.e., any row). With and without basal excitation is shown on the left and right columns respectively. **Top:** All cell populations. **Middle:** OLM cell LFP component. **Bottom:** PV cell LFP component.

From the LFP decompositions and different LFP patterns expressed (see Fig 7), and OLM cell activities (see Fig 2C), we can understand that the contribution of OLM cells is more dependent on *g_bs_* than *g_sb_* with the basal excitation affecting the peak power robustness more for larger *g_bs_* values. This is apparent in the color variation of the plots of the OLM cell LFP component in Fig 8 (middle). It is larger with basal excitation (left) than without basal excitation (right) for larger *g_bs_* values. Also, peak power magnitudes are larger with basal excitation. This is reflected in the mean and standard deviation without basal excitation (5.2 × 10^−10^, 2.2 × 10^−10^) that is smaller than with basal excitation (see values with basal excitation above). With only the PV cell LFP component, the LFP theta rhythm is disrupted as the interactions between basal excitation and PV inhibitory inputs are missing the OLM cell inputs. Specifically, the mean and std without basal excitation is (8.0 × 10^−9^, 1.1 × 10^−8^) which is smaller than with basal excitation (see values with basal excitation above). In essence, the inclusion of basal excitation can be considered as ‘adding’ to the magnitude and variance of the LFP power when OLM cells or PV cells are examined separately. In combination, a synergistic effect between inhibition and excitation occurs to generate a robust regime - a mean power with minimal variance. From Fig 2C, it can be seen that the PV cells (BC/AACs and BiCs) have activities that are more dependent on *g_sb_* than on *g_bs_*, and that BC/AACs are relatively more active than BiCs in the predicted regime. Thus, at larger *g_bs_* values when OLM cells are less active, BC/AACs would contribute more to keeping a synergistic balance with the basal excitation.

The LFP is generated on the basis of transmembrane currents. This means that the LFP is a weighted sum of inward and outward currents. How the LFP changes as a function of location is not trivial. When the LFP is governed by synaptic inputs the LFP peaks are narrower since the synaptic inputs are synchronized because of the coherent inhibitory spike rasters. On the other hand LFP signals governed by return currents would produce LFP peaks that are less narrow as the signal slows down as it travels down the dendrites producing a time lag. This all thus translates to synaptic input location dependencies. Thus, while we can visualize and appreciate the synergistic balances between excitation and inhibition from different cell populations, we note that these combinations are not easily seen as summated balances. Decompositions and intuitions from many simulations are required.

### LFP power across layers

As illustrated in Fig 6A, the color peak power plots are the power in the layer (particular electrode) where the power is maximal. To fully express this, we plot the maximum LFP power across the dendritic tree for all parameter sets in the predicted regime. This is shown in Fig 9 (left) with insets showing the same for the OLM cell (top) and PV cell (bottom) LFP components. From this, we see that the maximum LFP power is recorded at electrode 4, and that with only the OLM cell component, the power is distributed more widely and with only the PV cell component, more narrowly focused around the soma. This thus shows that the two populations differentially influence the location of LFP maxima. That the LFP power shows no discernible variability when all the cell populations are present, and that there is clear variability when not all of the cell populations are present is obvious in this Fig 9 (left). We did several additional sets of simulations to explore whether changes in the synaptic weights would affect whether the robust power feature in the predicted regime would still be present. In all the simulations presented so far, we used synaptic weights that did not bias the effect of one cell population type over the other based on their synaptic input location. So, for example, OLM cell inputs that are the furthest away from the soma had the largest synaptic weight. In doing this, we are following what was done previously in [19] who used ‘unbiased’ synaptic weights as well as using the same synaptic weight for all of the cell types. In using the same synaptic weight for all the cell types, we find that the robust power feature in the predicted regime remains (not shown).

**Fig 9.**
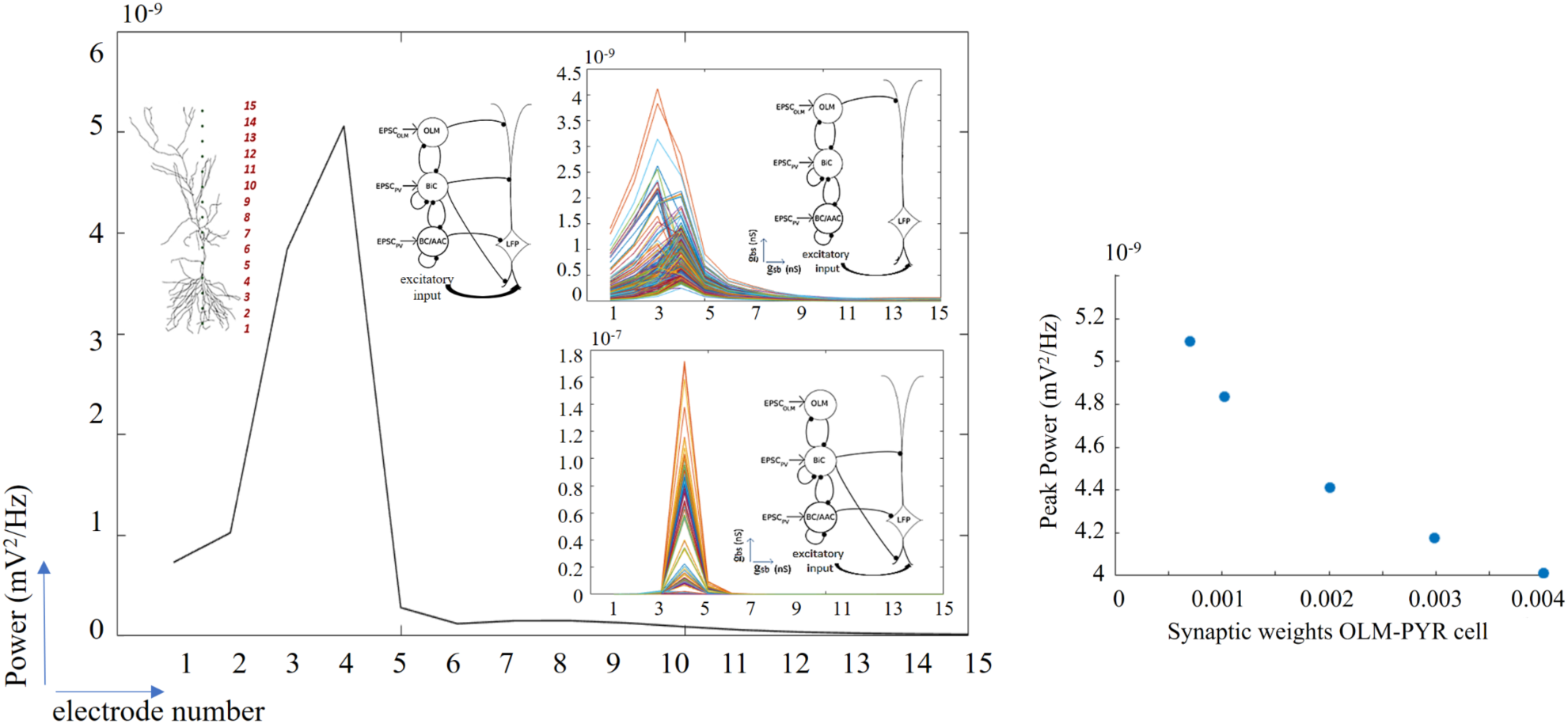
Laminar Power and Peak Power Changes With Changing Synaptic Weights. **Left:** Computed power at the different electrode locations to show laminar power distribution, for all sets of parameter values in the predicted regime. Top inset: Laminar power for OLM cell LFP component. Bottom inset: Laminar power for PV cell LFP component. Schematics shows the PYR cell model with the 15 extracellular electrodes and the different network con. **Right:** Changing the synaptic weight from the OLM cells to the PYR cell does not lead to much change in the peak power, as illustrated by the peak power at electrode 4. Parameter values: g_sb_ = 5.25, g_bs_ = 5.00 nS. Synaptic weights of 0.00067, 0.001, 0.002, 0.003, 0.004 µ S are shown.

As described and shown above, it is already clear that OLM cells via a direct OLM-PYR pathway minimally contribute to the LFP theta power. To show this directly, we did several, additional simulations where we changed the synaptic weight from OLM cells to the PYR cell. As an example, in Fig 9 (right) we show that increasing the synaptic weight by almost an order of magnitude decreases the peak power by about 20%.

### Estimating the number of PYR cells that contribute to the LFP signal

It is challenging to know how many cells contribute to an extracellular recording. The hippocampus has a regular cytoarchitecture with a nearly laminar, stratified structure of pyramidal cells [26]. This arrangement together with pyramidal cells being of similar morphologies and synaptic input profiles means that we can assume that any given pyramidal cell will generate a similar electric field leading to an additive effect with multiple cells in resulting LFP dipole recordings. Further, for the *in vitro* intrinsic theta LFP generation being considered in this work, the focus can be justified to the couple of synaptic pathways that we explored, and incoming inputs are synchronized amplifying the additive effect.

To estimate how many PYR cells contribute to an extracellular LFP recording in the *in vitro* whole hippocampus preparation, we define the ‘spatial reach’ of the LFP as the radius around the electrode where the LFP amplitude is decreased by 99%. Using our biophysical computational LFP models with parameter values taken from the predicted regime, we find that the spatial reach is 300 *µ*m as measured extracellularly close to the soma since the LFP decreases from 10,000 nV to 100 nV within this radius. This is shown in Fig 10 where the dotted arrow represents this radius. The spatial reach estimate is done using an electrode placed extracellularly close to the soma of the pyramidal cell, in stratum pyramidale, since this is where maximal power is recorded (see Fig 9). Taking advantage of detailed quantitative assessment and modeling done by Bezaire and colleagues [25, 27], there are about 311,500 PYR cells in a volume of 0.2 mm^3^ of ‘stratum pyramidale’ tissue (see model specifics in Fig 1 of [27]). Given our spatial reach radius estimate, a cylindrical volume of stratum pyramidale would be 0.014 mm^3^ or about 7% of the total number of PYR cells which is about 22,000. In this way we estimate that there are about 22,000 PYR cells that contribute to the LFP signal. We note that this is an upper bound, as we assume correlated activity across pyramidal cells and homogeneous extracellular electrical properties. The presence of membranes and biological compounds in the extracellular space will in principal decrease the propagation of the LFP.

**Fig 10.**
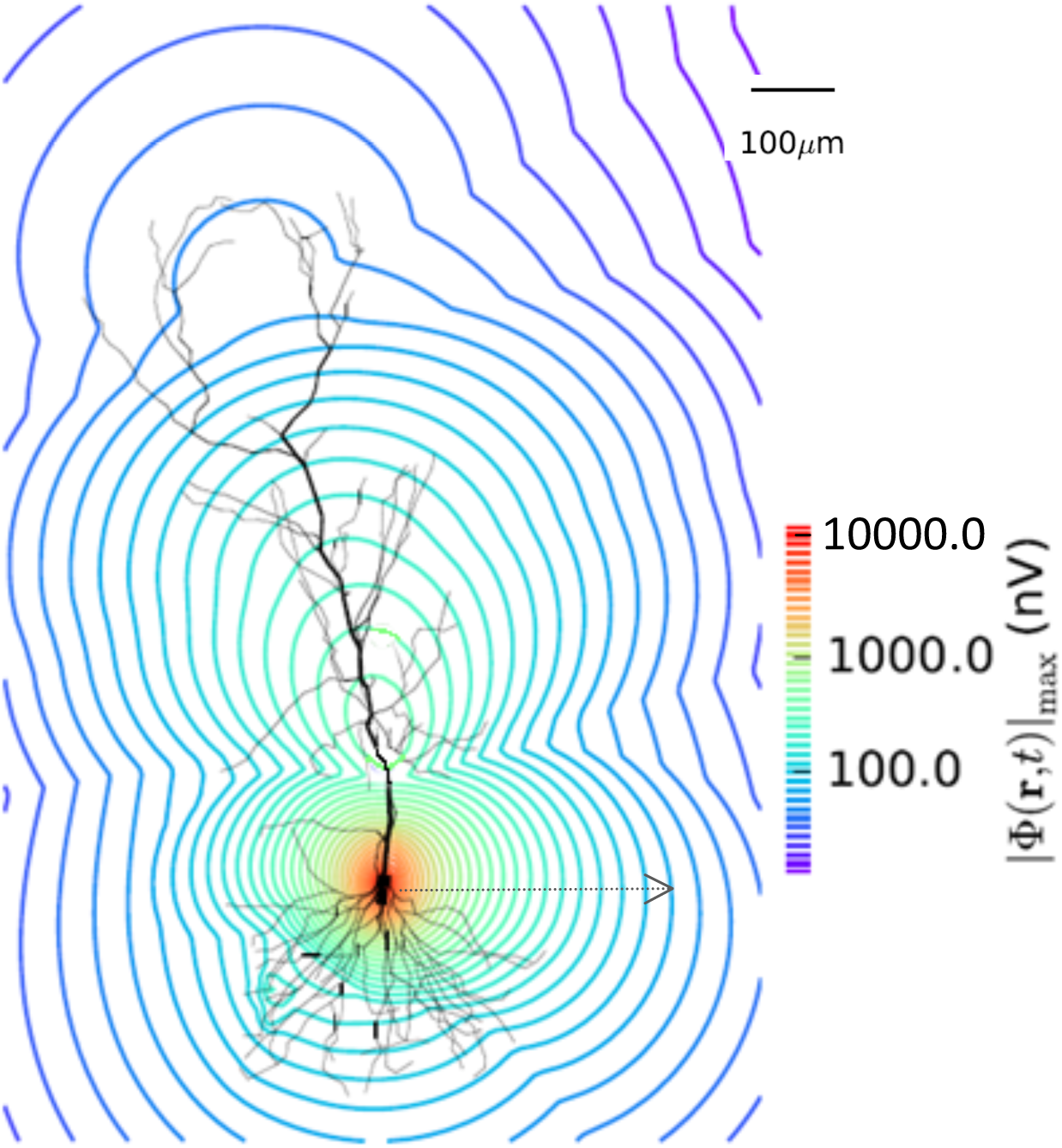
Spatial Attenuation. We estimate the size of the region an LFP electrode can ‘see’ using our models. PYR cell model morphology is shown with calculated signal decrease from a electrode positioned near the cell soma. The dotted arrow shows the extent of the spatial reach of the signal that is taken as a 99% decrease in the signal, and is approximately 300 *µ*m. Parameter values used are from the predicted regime. g_sb_=5, g_bs_=5.75 nS, c_sb_=0.21.

## Discussion

To a large extent, understanding brain function and coding requires that we are able to understand how oscillatory LFP signals are generated [3, 28, 29]. Cross-frequency coupling analyses of LFP signals has led to ideas underlying learning and memory functioning [30], and it is always important to do careful analyses [31]. Further, given that particular inhibitory cell populations and abnormalities in theta rhythms are associated with disease states [8], we need to consider how different cell types and pathways contribute to LFP recordings. Ultimately, the challenge is to bring together LFP studies from experimental, modeling and analysis perspectives. In this work, we make steps toward this challenge by gaining insight into the contribution of OLM cells to theta rhythms. We took advantage of an *in vitro* whole hippocampus preparation that spontaneously expresses *intrinsic* theta rhythms [15], and our previous inhibitory network models developed for this experimental context [19], to build biophysical LFP models. We leveraged our LFP models to make direct comparison with experimental LFP characteristics. This allowed us to predict coupling parameters which in turn led us to determine OLM cells’ contribution to intrinsic theta LFP rhythms.

We showed how the extracellular theta field recorded along the cellular axis of a PYR cell is affected by the magnitude of the inhibitory synaptic currents inserted along its dendritic arbor. Fluctuations in the magnitude of the total inhibitory input occur due to alterations in synaptic strength balances of the inhibitory networks. Our models exhibited network states in which interactions between OLM cells and BiCs could invert the polarity of the recorded signal and produce extracellular potentials of high or low magnitude. We also distinguished regimes where these cellular interactions preserved the frequency of the signal versus those that led to lags or abolishments of the extracellular LFP rhythm. When we applied experimental characteristics of theta frequencies and polarities to our biophysical LFP models, a clear selection emerged and thus we were able to predict parameter values regarding connectivities. Specifically, we found that the connection probability from OLM cells to BiCs is 0.21 and that synaptic conductances from OLM cells to BiCs had to be larger than 3.5 nS, and we called this the *predicted* regime.

Unexpectedly, we found that this predicted regime also exhibits a robust power output. That is, so long as parameter values were within the predicted regime, the power did not change (Fig 6A), and in this regime we see that BiCs are mostly silenced, BC/ACCs are significantly active while OLM cell activity decreases from high to low values as *g_bs_* increases (Fig 2C). By decomposing the signal we revealed that OLM cell inputs minimally contributed to the LFP power unlike the other cell populations (BiCs and BC/AACs or PV cells). The power of the OLM cell LFP component on its own, although low, showed some variation in the predicted regime (coefficient of variation or CV < 1). On the other hand, the power of the PV cell LFP component was a couple of orders of magnitude higher and showed more variation (CV > 1) in the predicted regime. This indicates that OLM cells contribute to LFP power robustness without contributing to average power whereas PV cells contribute to average power but their effect is more sensitive to perturbations in OLM-BiC interactions. Therefore their contribution is variable. It is however interesting to note that the PV LFP component average power was larger than the average power of the predicted regime with all cells being present. Thus our results indicate that adding OLM cells in the network can overall cause a small decrease in LFP average power as compared to when only PV cells are present and of course induce robustness. It was also interesting to observe that in almost half of the cases the OLM cell LFP component was arrhythmic or non-oscillatory despite the fact that OLM cells are driven by theta paced EPSCs. That is, OLM cell inputs alone in most cases are not able to generate a theta LFP signal as recorded in the extracellular space of the PYR cell even though OLM cell populations themselves are firing at theta frequency. Further LFP signal analysis decomposition showed that removing only basal excitation disrupted the robustness of the predicted regime. This suggests that a synergy of OLM cell inputs and basal excitatory inputs as co-activation of distal inhibition and proximal excitation is important to produce robustness in the predicted regime. Overall, an essential aspect in comparing model and experiment LFPs to predict model parameters and decipher cellular contributions was to match sources and sinks at different layers. Thus, having recordings from multiple layers is important.

### Morphological details, synaptic locations and related studies

As the main contribution to the LFP is thought to stem from synaptic input to neurons and the subthreshold dendritic processing, various studies have investigated how morphological characteristics and intrinsic resonances shape the features of the LFP signal. In most cases input synapses are activated according to Poissonian statistics [32–34]. However, in our study here the origin population consisted of point neuron cell representations that had been constrained based on experimental patch clamp recordings from the whole hippocampus preparation. Essentially, we have used a hybrid scheme which is a combination of a point neuron network origin population contacting a multi-compartment PYR cell model which serves as a processor of synaptic inputs and produces the LFP.

One factor modulating the amplitude of LFPs is related to the somatodendritic location of synaptic inputs on the PYR cell tree. Different populations of GABAergic interneurons target different dendritic domains and the domain-specific targeting of various interneurons supports the hypothesis of domain-specific synaptic integration in CA1 PYR cells [35]. In CA1 PYR cells, distal and middle apical dendrites comprise two distinct dendritic domains with separate branching connected by a thick apical dendrite. This cytoarchitectonic separation of the cluster of distal dendrites relative to middle and proximal dendrites was shown to critically reduce the effect of distal EPSCs to somatic excitability [36]. The presence of a single apical dendrite with many obliques in stratum radiatum caused a large shunting of EPSCs traveling from the tuft dendrites to the soma. Thus we can appreciate our observation that OLM cells, which target distal dendrites, minimally affected LFP power in stratum pyramidale considering the limited ability of distal inhibition to reach more proximal and somatic regions of the CA1 PYR where maximum power was recorded. This is not just due to the distal location of these inputs but more due to the cytoarchitectonic separation of the cluster of distal dendrites relative to middle and proximal dendrites. This separation prohibits inhibitory inputs in distal regions from effectively propagating to somatic and proximal regions of CA1 PYR cells and thus being reflected in the extracellular space.

We can further consider our results in light of another theoretical modeling study by Gidon and Segev [37] which showed that inhibitory inputs can affect excitatory inputs locally and/or globally, depending on the relative locations of the excitatory and inhibitory synapses. In particular this can help us understand the loss of robust power in the predicted regime after removal of OLM cells. The predicted regime consists of different connectivities that generate different spiking patterns that give rise to fluctuations in inhibitory input in different synaptic locations. First, inhibitory input hyperpolarizes the membrane potential, which results in shunting of the adjacent dendritic compartments. Activation of excitatory synapses within the shunted compartments will thus generate smaller depolarization, compared with non-shunted dendrites (“local” effect). Second, the local shunting would suppress excitatory input in a nonlinear fashion at locations that are not directly affected by the shunting (“global” effect). Thus, when inhibitory inputs are activated simultaneously with excitatory inputs, the average (i.e., across trials) evoked membrane potential within shunted dendritic compartments should be smaller compared with compartments that have no inhibitory input. At the same time, excitatory effects throughout the entire dendritic tree would be reduced in a nonlinear fashion, and which can be quantified as the change (with versus without inhibitory input) of the trial-to-trial variability of the membrane potential. In our case the activation of excitatory inputs occurs in regions not close by the OLM cell inhibitory inputs, thus the overall power does not increase but the robustness is affected. In [37] the authors examined the spread of shunt level implications using a CA1 reconstructed neuron model receiving inhibition at three distinct dendritic subdomains: the basal, the apical, and the oblique dendrites as innervated by inhibitory synapses. They found that the shunt level spread effectively hundreds of micrometers centripetally to the contact sites themselves spanning from the distal dendrites to the somatic area. This observation thus shows that the somatic area is indeed influenced by shunting inhibition which means that excitatory input non-linearities in our model will be reduced in the presence of global inhibition in the somatic area leading to a decrease in variability and thus robustness in the membrane potential. Of course, the LFP is a measurement of transmembrane currents and not membrane potential. However the reduction of excitatory input mediated non-linearities will also reduce the variability in the distribution of return currents and thus the variability in the LFP.

### Limitations and future considerations

Our present study was limited in terms of not considering more inhibitory cell types (e.g., see [27]) and by considering *ongoing* intrinsic theta rhythms since theta frequency inputs were used (Fig 1). However, our inhibitory network models were constrained by the experimental context and our less complex model representations enabled us to explore many thousands of simulations and directly compare our biophysical LFPs with experimental LFP features. This aspect was key in allowing us to predict parameter value sets and gain insights.

Theta rhythms are foremost generated due to subthreshold activity and dendritic processing of synaptic inputs. Here we used a passive PYR cell model as the spiking component has been shown to mainly contribute to the LFP at frequencies higher than 90Hz [6] while the active voltage-gated channels that have been eliminated here have also been shown to influence LFP characteristics more prominently in frequencies above the theta range [38]. Thus, although the presence of voltage-gated channels will influence the exact distribution of return currents, we thought that it was a reasonable simplification to not include them in this study.

Extracellular studies suggest that the main current generators of field theta waves are the coherent dendritic and somatic membrane potential fluctuations of the orderly aligned pyramidal cells [39–41]. Thus, distal and local ascending pathways onto PYR cells can in principle contribute to extracellular LFP deflections. To understand theta rhythms one needs to consider the populations projecting onto the PYR cells in CA1. During *in vivo* behaviors, medial septum and entorhinal cortical inputs onto CA1 PYR cells are prominent modulators of the amplitude, phase and waveform features of theta rhythms in conjunction with local inhibitory and excitatory cells. However, spatiotemporal coincidence of inputs makes separation difficult and thus it is challenging to determine cellular contributions to LFP recordings. As there is significant spatiotemporal overlap on PYR cell dendrites across ascending pathways it would be hard to disentangle the cellular composition of these pathways and assess the cellular contribution to theta LFP characteristics. As shown in previous studies [42] blind separation techniques such as Independent Component Analysis (ICA) produce poor results when trying to disentangle combinations of rhythmic synaptic sources with extensive spatiotemporal overlap. By focusing on intrinsic theta rhythms in the *in vitro* whole hippocampus preparation here, we could reduce the spatiotemporal overlap of different pathways and unravel the cellular composition of the different pathways projecting to the PYR cell. We were thus able to decipher the contribution of OLM cells to intrinsic theta rhythms. This work could potentially be used as a basis to understand OLM cell contributions during *in vivo* theta LFP recordings.

Moving forward we aim to take advantage of the insights gained here to build hypothesis-driven theta generating networks. In this way, we hope to be able to determine the contribution of different cell types and pathways to LFP recordings that are so heavily used and interpreted in neuroscience today.

## Methods

### Network model details

This work builds on previously developed models described in [19]. Here we provide a summary of specifics that are salient to the present study.

#### Inhibitory cell types and numbers, PYR cell model

The inhibitory network model consists of 850 cells that represents a volume of 1 mm^3^ as shown to be appropriate to obtain spontaneous theta rhythms in the *in vitro* whole hippocampus preparation [15, 19, 20]. Four different types of inhibitory cells are included: basket/axo-axonic cells (BC/AACs), bistratified cells (BiCs) and oriens-lacunosum/moleculare (OLM) cells. BC/AACs comprise a 380-cell population and target somatic, perisomatic and axo-axonic regions of pyramidal (PYR) cells. The BiCs comprise a 120-cell population and target middle, apical and basal regions of PYR cells, and the OLM cells comprise a 350-cell population and target the distal, apical dendrites of PYR cells. As in [19], the structure of the PYR cell model is based on the one used in [43] as implemented in the NEURON Simulator [44] (see ModelDB Accession number 144541). The PYR cell model is used as a passive integrator of inputs from cell firings at the various layers of the hippocampus, and all active, voltage-gated channel conductances are set to zero. This overall network model is schematized at the top of Fig 1. With the exception of basal excitatory input, it is the same as used in [19]).

#### Inhibitory cell models and drives

The inhibitory cell models are single compartment, have an Izhikevich mathematical structure [45] and were constructed by fitting to experimental data from whole cell patch clamp recordings in the whole hippocampus preparation [19]. All of the cell model parameter values are given in [19]. PV cell types are BC/AACs and BiCs, and SOM cell types are OLM cells. Each cell model is driven by excitatory postsynaptic currents (EPSCs) taken directly from experiment [23] during ongoing spontaneous theta rhythms for PV or SOM cells. As detailed in [19], we designed the EPSCs to ensure that the inhibitory cells received frequency-matched current inputs and at the same time had amplitudes and peak alignments that were consistent with experiment (EPSC_*PV*_ and EPSC_*OLM*_ examples in Fig 1 top). Importantly, we captured the experimental variability in amplitude and timing of EPSCs across cells by varying the gain (factor by which the EPSC was scaled to alter the amplitude) and timing of the EPSCs across cells with a normal distribution in accordance with the experimental recordings. Thus, each inhibitory cell model received a unique set of excitatory synaptic inputs reflecting the range of amplitudes and timing of those recorded experimentally.

#### Inhibitory network connectivity and output

PV cells (BC/AACs and BiCs) were randomly connected with probabilities and synaptic conductance values based on experimental estimates from the literature and our previous modeling work [20]. Connections between BiCs and OLM cells are known to exist [17] and we previously estimated a range of values from the literature, with the connection probability from BiCs to OLM cells taken as 0.64 times the connection probability from OLM cells to BiCs [19]. Also from the literature, we estimated synaptic conductances between OLM cell and BiCs. Although OLM-BiC connections exist, their values are unclear and in our previous work we specifically examined what sort of balance of parameter values would be important in theta rhythms by examining a wide range of values that encompass our estimates [19]. Inhibitory synapses were modeled using a first order kinetic process with appropriate rise and decay time constants. For the work in this paper, we use the output from the inhibitory networks. Thus, inhibitory connectivities used here are the same as in [19] and are shown in Table 1. The spiking output of the inhibitory network models briefly described above were computed for the range of synaptic conductance strengths and connection probabilities as given in Table 1. Specifically, these simulations were done for 5 seconds; the connection probability from OLM cells to BiCs (*c_sb_*) varied from 0.01 to 0.33 with a step size of 0.02 producing 16 sets of connection probabilities; synaptic conductance values range from 0-6 nS for OLM cells to BiCs (*g_sb_*) and for BiCs to OLM cells (*g_bs_*). By changing *g_sb_* and *g_bs_* with a step size resolution of 0.25 nS, 625 raster plots are produced. So the total number of raster plots used in our study here as computed in [19] is (625 × 16) 10,000, and they are all available on Open Science Framework (osf.io/vw3jh).

#### Synaptic weights and distribution onto PYR cell

Inhibitory inputs to the PYR cell model were distributed in the same way as done in [19]. That is, we distinguished between synapses at the distal layer (stratum lacunosum-moleculare), medial and basal layers (stratum radiatum and oriens), and the perisomatic/somatic layer (stratum pyramidale). Distal synapses were defined as those that were > 475*µ*m from the soma; apical and basal synapses are defined as those that were 50 − 375*µ*m from the soma; perisomatic/somatic synapses are defined as those which were < 30*µ*m from the soma. We created three lists of components (where each component points to a specific segment of a section in the PYR cell model), for the possible distal, proximal apical/basal, and perisomatic/somatic synaptic targets. For each individual, presynaptic inhibitory cell model, we randomly chose a synaptic location on the passive CA1 PYR cell model from the respective list (distal dendrites for OLM cell models, apical/basal dendrites for BiC models, and perisomatic/somatic locations for BC/AACs). Then the spike times from the individual, inhibitory cell models filled a vector, and an artificial spiking cell was defined to generate spike events at the times stored in that vector at the specific location at which that cell creates a synaptic target. We used the Exp2Syn function in NEURON to define the synaptic kinetic scheme of the synapse. This function defines a synapse as a synaptic event with exponential rise and decay, that is triggered by presynaptic spikes, and has a specific weight that determines its synaptic strength, and an inhibitory reversal potential of −85 mV, as measured in the whole hippocampus preparation. Synaptic weight values onto the PYR cell from the different cell populations were estimated using somatic IPSC values for OLM cells onto PYR cells [46]. As these synaptic weights are not clearly known, we used two different synaptic weight profiles in the explorations of our previous work [19]. One profile in which the synaptic weights were all the same (0.00067 *µ*S) as estimated from the OLM cells IPSC currents, and another profile that was graded such that the different cell types led to similar somatic IPSC amplitudes. Finally, we note that in our previous work [19] we used an ad-hoc representation for LFPs which was an inverted summation of all integrated inputs as measured at the PYR cell soma. That is, the postsynaptic potentials on the PYR cell are due to the various inhibitory cell firings that comprise the presynaptic spike populations.

### Additional network model details for this study

For the study here, inhibitory inputs were distributed in the same way as in [19], and we used graded synaptic weights for the bulk of our simulations - these values are shown in Table 1. However, other values were also explored. In [19] we used the literature to estimate that synaptic conductances between OLM cells and BiCs are 3-4 nS, and Bezaire et al [27] used 10 synapses/connection as estimates in their detailed data-driven computational models. This implies that a single synapse would be 0.3-0.4 nS, representing an approximate minimum connection weight.

As direct comparisons were made with theta LFP experimental recordings, it was important to include excitatory input to the PYR cell model. Thus we also included excitation due to CA1 recurrent collaterals which synapse on basal dendrites [24]. As noted in [19], excitatory feedback was not included in a direct fashion as we were focused on ongoing theta rhythms and OLM-BiC interactions, and not theta generation mechanisms explicitly, and we did not explicitly model excitatory cell populations as we did for inhibitory cell populations. This means that we do not have spike rasters for excitatory populations as we do for the inhibitory cell populations. Rather than generate an arbitrary set of spike times to simulate excitatory inputs, we used spike times from a BiC raster (*g_sb_*=3.75, *g_bs_*=1.75 nS) in which the neuron order was randomized, and comparable synaptic weights were used. Using these random spike trains we generated spike vectors exactly as in the case of interneurons and randomly distributed them on basal dendrites using 197 synapses based on number estimates from [25, 27]. In this way, we did not have a spatiotemporal dominance of inhibitory or excitatory input in basal dendrites. We used an excitatory reversal potential of −15 mV as measured in the whole hippocampus preparation, and synaptic time constants in line with our modeling work [47]. In essence, we simulated EPSCs using random spike trains of theta frequency instead of explicitly modeling pyramidal cell spiking activity. We note that with these choices, somatically recorded currents in our PYR cell models are similar to what is observed in experiments [23]. All parameter values are summarized in Table 1.

We note that the inhibitory cell spike rasters used from our previous study [19] used random connectivities between the different inhibitory cell populations. Let us consider that a given set of parameters (*c_sb_*, *g_sb_*, *g_bs_*) defines a connectivity map. Each cell within a given population is randomly assigned a synaptic location within the boundaries of the dendritic tree on which it projects. Based on a given connectivity map the spiking activity of the various cell populations will differ. Therefore the characteristics of the produced biophysical LFP will depend on the spike distribution of a given population defined by the connectivity map and also the number and location of synapses on the dendritic tree. To ensure that our LFP output was not dependent on the specific synaptic location that every cell was assigned to, we generalized our observations by performing many trials for a given connectivity map (that had a given connection probability and synaptic conductance values), assigning randomly different location to the cells of each population to ensure that the LFP output was not dependent on that aspect. The effect of not having the same, exact spatial location of synapses can be specifically observed when our LFP model was decomposed to only have OLM cell inputs (Fig 6B). Here, when *g_bs_* = 0, the LFP power should be the same, but it is not because the spatial location of OLM cell to PYR cell synapses are not identical. Since the overall power is much less than when only PV cell inputs are present and there is alot of space for currents to exit, the non-exact spatial location of OLM cell synapses is apparent in the resultant peak power not being exactly the same when *g_bs_* = 0. This is unlike the case when only PV cells inputs are present (Fig 6C) and when *g_sb_* = 0. The LFP power is the same even though the spatial locations for PV cell synapses are not exactly the same. Here, there are more inputs in a smaller spatial region so that not having the exact same spatial location does not have observable effects on the peak power.

### Biophysical computation of LFP

Extracellular potentials are generated by transmembrane currents [48]. In the commonly used volume conductor theory, also used here, the extracellular medium is modeled as a smooth three-dimensional continuum with transmembrane currents representing volume current sources. The fundamental formula relating neural activity in an infinite volume conductor to the generation of the LFP w(t) at a position r is given by [18].

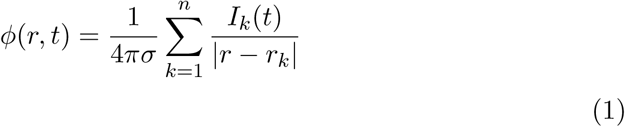

Here I_k_ denotes the transmembrane current (including the capacitive current) in a neural compartment *k* positioned at *r_k_*, and the extracellular conductivity, here assumed real (ohmic), isotropic (same in all directions) and homogeneous (same at all positions), is denoted by σ. In the hippocampus the mean extracellular conductivity σ is equal to 0.3Sm^−1^ [49] which is the value that we have used for our simulations. A key feature of Equation 1 is that it is linear, i.e., the contributions to the LFP from the various compartments in a neuron sum up. Likewise the contributions from all the neurons in a population add up linearly. The transmembrane currents *I_k_* setting up the extracellular potentials according to Equation 1 are calculated by means of standard multi-compartment modeling techniques, here by use of the simulation tool NEURON [44]. The current source densities (CSDs) in Fig 1 were computed using the 1D kCSD inverse method proposed in [50]. The CSDs are computed from the LFP measured by electrodes that are arranged along a straight line, in this case along the cellular axis of the PYR cell.

The same PYR cell multi-compartment model as described above was used to compute the extracellular biophysical LFP, and we used the set of 10,000 5-second raster plots (of inhibitory spikes) as generated previously for our presynaptic populations with the addition of basal excitation. That is, we generate extracellular potential traces (5 sec each) due to the various inhibitory cell firings (10,000 raster plot sets).

### Simulation details

The computational simulations and analyses are performed using the LFPy python package [33], NEURON [44], and “MATLAB 2015” [51]. The large scale network simulations were conducted using high-performance computing at SciNet [52]. Code is available on https://github.com/FKSkinnerLab/LFP_microcircuit.

## Acknowledgments

We thank Katie Ferguson for her feedback and participation at early stages of this work, Benedicte Amilhon and Sylvain Williams for their helpful comments, and Alexandre Guet-McCreight for a careful reading of this work. We thank NSERC Canada, Margaret J. Santalo Scholarship (Dept Physiology, Toronto), OGS and the SciNet HPC Consortium for support.

## Supporting information

**S1 Fig.**
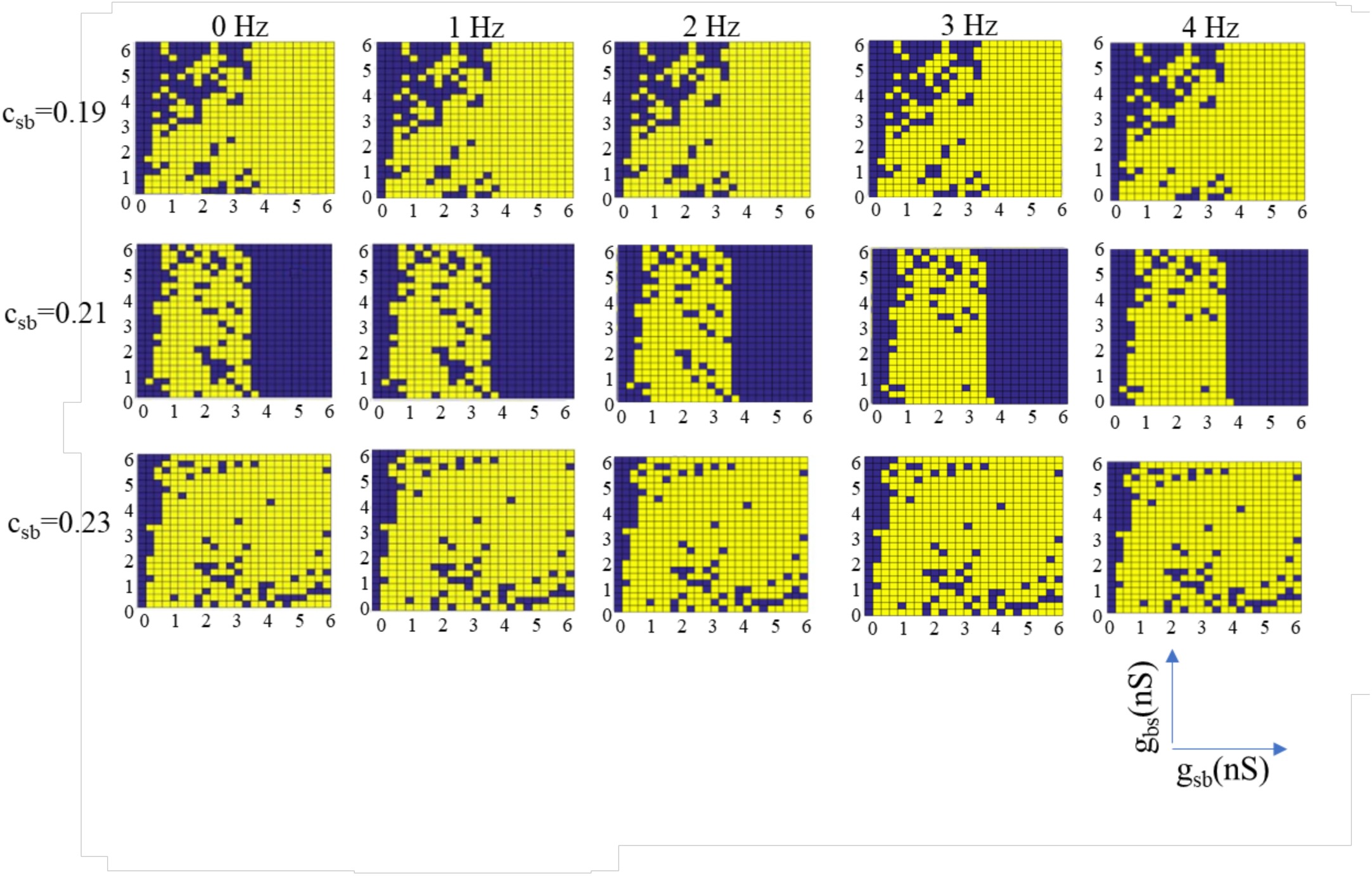
Selected and Rejected Parameter Sets Using Different Frequency Bounds.

**S2 Fig.**
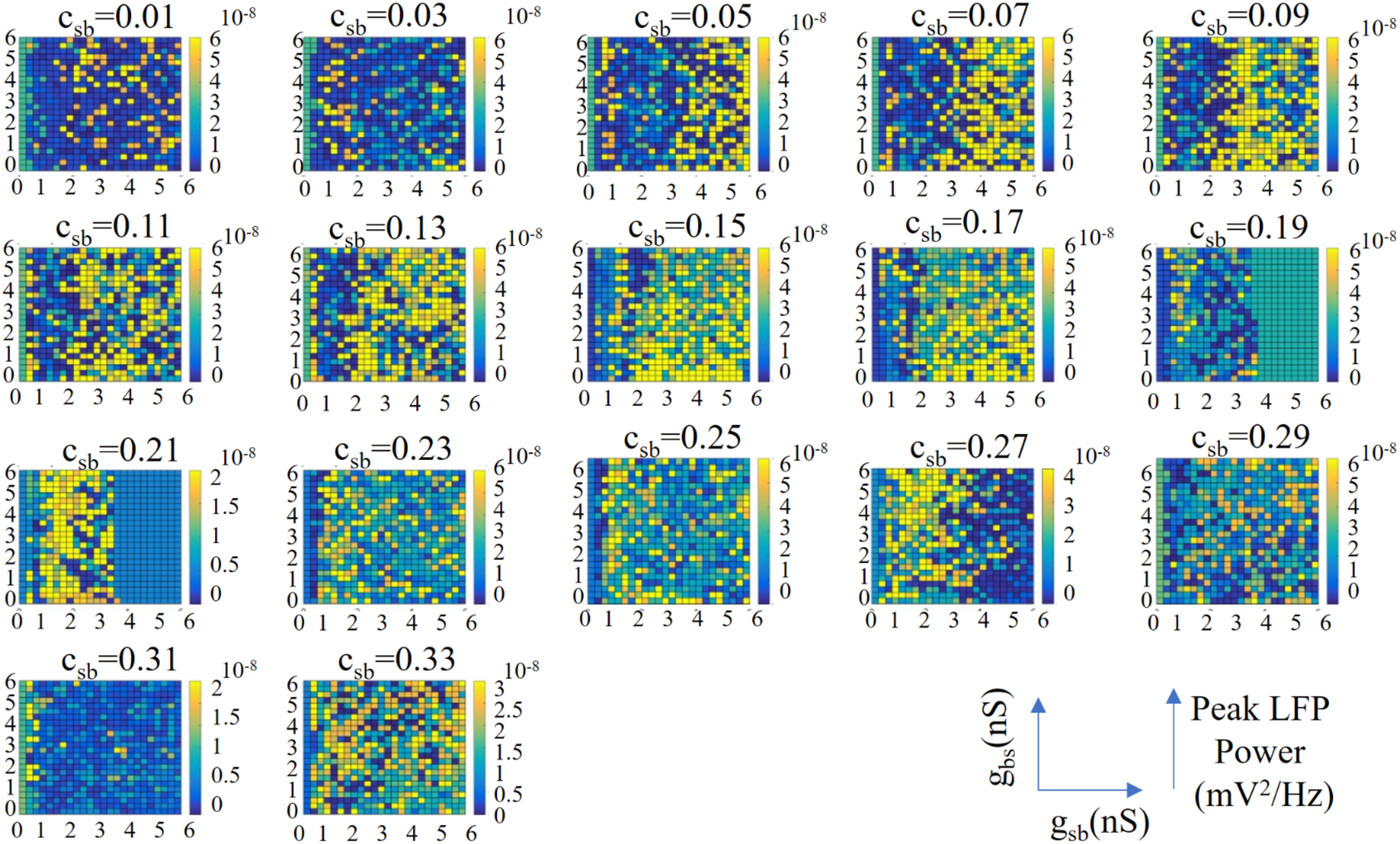
Peak Power For All Conductances and Connectivities.

